# Ablation of *Prdm16* and beige fat causes vascular remodeling and elevated blood pressure

**DOI:** 10.1101/2025.06.12.658904

**Authors:** Mascha Koenen, Tobias Becher, Giulia Pagano, Ilaria Del Gaudio, Jorge A Barrero, Augusto C Montezano, Jenelys Ruiz Ortiz, Zeran Lin, Nicolás Gómez-Banoy, Rose Amblard, Meltem E Kars, Luisa Rubinelli, Sarah J Halix, Zhen Fang Huang Cao, Xing Zeng, Scott D Butler, Yuval Itan, Rhian M Touyz, Annarita Di Lorenzo, Paul Cohen

## Abstract

While excess adiposity is a major risk factor for hypertension and cardiovascular disease, brown fat is associated with protection from these pathologies. Whether brown fat has a causal role in this process and the underlying molecular mechanisms remain unknown. Here we investigate the role of murine beige fat, as a model of inducible brown fat in humans, in adipocyte-vascular crosstalk. Using mice with an adipocyte-specific deletion of PRDM16, resulting in a loss of beige adipocyte identity, we discover a dramatic remodeling of perivascular adipose tissue, increased vascular reactivity and elevated blood pressure. We further show that the circulating enzyme *Qsox1* is de-repressed in *Prdm16*-deficient adipocytes, and deletion of *Qsox1* in PRDM16cKO mice rescues vascular fibrosis and reactivity. These results demonstrate a key new role for beige adipocytes in blood pressure regulation and identify *Qsox1* as an important mediator of adipocyte-vascular crosstalk.

## INTRODUCTION

Hypertension is a significant risk factor for coronary heart disease and stroke (1,2), and even mild, chronically increased blood pressure is associated with end-organ damage and increased mortality (3–7). Obesity is a major risk factor for hypertension, though the underlying mechanisms are not fully understood. Adipose tissue crosstalk with the vasculature is an important determinant of blood pressure regulation with both loss (8,9) and excess accumulation (10,11) of adipose tissue being associated with detrimental phenotypes. The type of adipose tissue (12–14) more than the total amount is a particularly critical factor in blood pressure regulation. Whereas excess white fat, especially visceral adiposity, is associated with elevated blood pressure (10), brown fat is associated with reduced odds of hypertension, even in obesity (15).

Brown adipose tissue is best known for its ability to produce heat (thermogenesis) through uncoupled respiration, but is increasingly recognized as an endocrine organ (16–18). There are two types of thermogenic brown fat: developmentally preformed classical brown fat (BAT) and inducible beige fat (19). Comparative transcriptomic analyses suggest that inducible thermogenic fat in adult humans is more similar to murine beige fat (20–22), making it an important target to understand adipose-to-vascular crosstalk. In mice, beige adipocyte biogenesis and function are dependent on the transcriptional coregulatory protein PR domain-containing 16 (PRDM16) (23). Ablation of *Prdm16* in adipocytes (PRDM16cKO) leads to the loss of beige fat identity (23), providing a unique genetic tool to study this tissue’s impact on vascular function.

Humans and mice possess multiple thermogenic fat depots (19), including perivascular adipose tissue (PVAT) surrounding the thoracic aorta and within the carotid sheath (24–27). Remodeling of PVAT during obesity or aging (28–32) and genetic ablation result in accelerated inflammation and hypertension (9,33), supporting a cardioprotective role for healthy PVAT. The contribution of beige adipocytes to this protective function, however, has yet to be investigated. Here, we used PRDM16cKO mice to ablate beige adipocytes and investigated the relationship between thermogenic adipose tissue, vascular remodeling and blood pressure regulation, elucidating a new adipocyte-vascular crosstalk explaining the cardiovascular benefits of thermogenic fat observed in humans.

## RESULTS

### PRDM16-dependent adipocyte remodeling regulates vascular function

Aortic PVAT is comprised of different types of adipocytes (34), with the thoracic aortic PVAT (tPVAT) predominantly containing multilocular brown/beige adipocytes (Fig. 1A) (25,27,35) and the abdominal aortic PVAT (aPVAT) mostly consisting of unilocular white adipocytes (36,37) and only a few multilocular adipocytes (Fig. S1A, black arrows). In addition to previously described changes in subcutaneous fat (23), we found that adipocyte-specific deletion of *Prdm16* (PRDM16cKO) results in drastic remodeling in PVAT (Fig. 1B and Fig S1B, red arrow). Along with a loss of multilocular lipid droplets, uncoupling protein 1 (UCP1), a marker of thermogenic adipocytes, was downregulated in PRDM16cKO PVAT (Fig 1C, D and Fig S1C, D), whereas the cold-stimulated induction of UCP1 was not affected (Fig 1E, F). RNA sequencing of both PVAT depots (Fig 1G, Fig S1E-H) demonstrated widespread loss of the thermogenic signature (*Ucp1, Prdm16, Cpn2, Cox8b*) (Fig 1G, Fig S1H) and a concomitant increase in white adipocyte identity markers (*Aldh3b2, Reep6, Agt*) in PRDM16cKO mice (Fig 1G, L, Fig S1H), which was further corroborated at the protein level (Fig 1H-K and Fig S1K). Pathway enrichment analysis supported these results (Fig S1I, J), illustrating a previously undescribed PRDM16-dependent remodeling of PVAT, independent of changes in body weight (Fig 1M). The dramatic increase in angiotensinogen (AGT) in tPVAT of PRDM16cKO mice (Fig 1G, J-L) was particularly noteworthy, as AGT is the precursor of the potent vasoconstrictor angiotensin II (ANGII) and is directly transcriptionally repressed by PRDM16 (38,39). Previous work has shown that local adipocyte-derived AGT signaling can increase circulating AGT (40,41) however, we did not find increased circulating levels of AGT (Fig 1N) and ANGII (Fig 1O) in PRDM16cKO mice.

**Fig 1.**
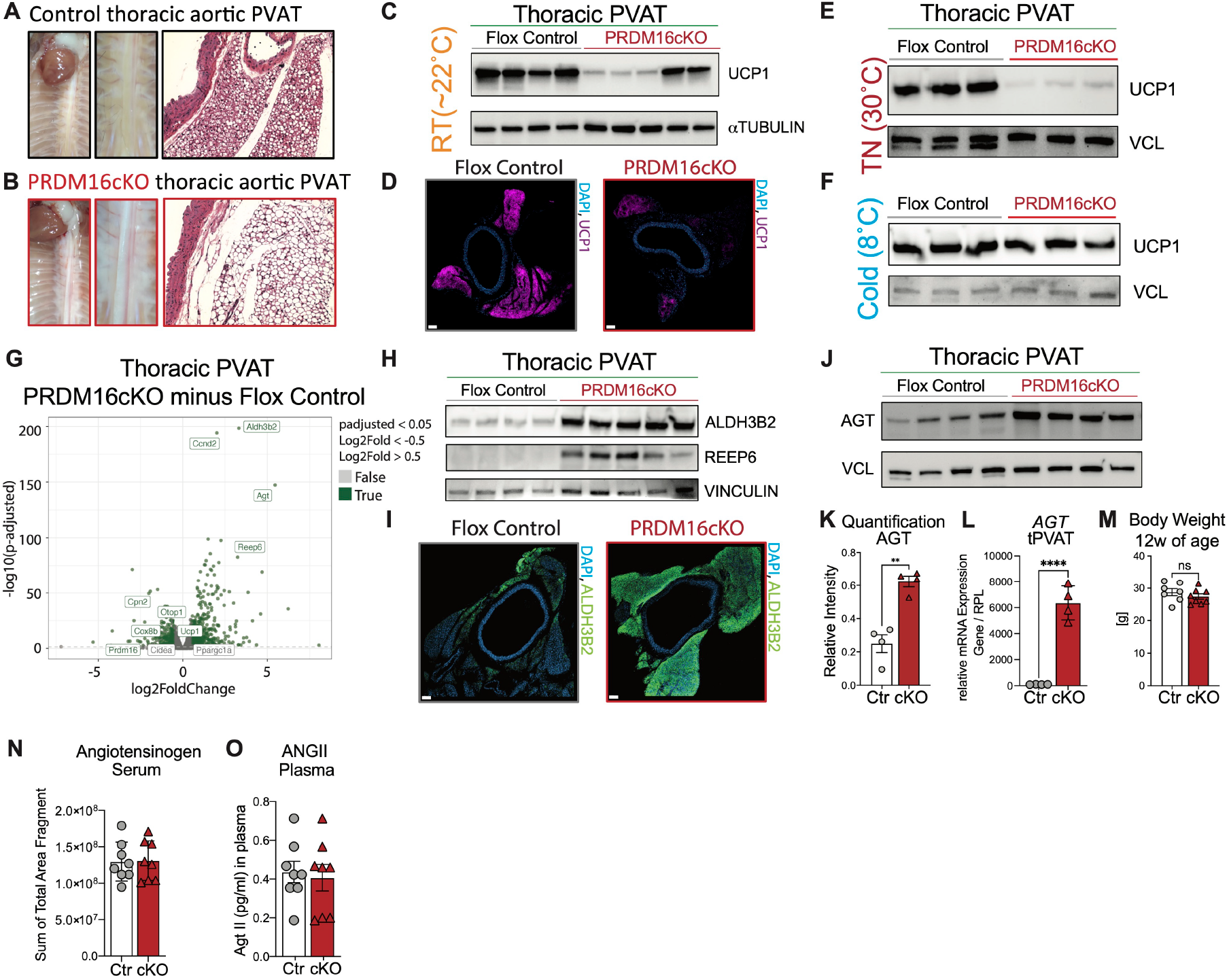
Loss of *Prdm16* in adipocytes leads to drastic remodeling of thoracic aortic PVAT. **(A, B)** Representative appearance and HE stain of the thoracic perivascular adipose tissue (PVAT) in (A) flox control and (B) PRDM16cKO mice. (**C**) Representative Western Blots of UCP1 and αTUBULIN of flox control (n=4) and PRDM16cKO (n=5) tPVAT and (**D**) Representative immunofluorescence images of UCP1 and DAPI staining on flox control (grey) and PRDM16cKO (red) tPVAT. (**E, F**) Representative Western Blots of UCP1 and VINCULIN of flox control (n=3) and PRDM16cKO (n=3) housed at (E) thermoneutrality (TN, 30°C) or (F) cold (8°C) for 1 week. (**G**) Volcano plot for DEGs in thoracic PVAT of PRDM16cKO minus flox control (green color = significantly up-/down-regulated genes (padj. <0.05) and |log_2_[fold change] | ≥ 0.5 and grey color display non-significant DEGs). Some significant DEGs related to thermogenesis and most upregulated in PRDM16cKO are highlighted. (**H**) Representative Western Blots of ALDH3B2, REEP6 and VINCULIN of flox control (n=4) and PRDM16cKO (n=5) tPVAT and (**I**) Representative immunofluorescence images of ALDH3B2 and DAPI staining on flox control (grey) and PRDM16cKO (red) tPVAT. (**J**) Representative Western Blots of AGT (Angiotensinogen) of flox control (n=4) and PRDM16cKO (n=5) tPVAT and (**K**) quantification. (**L**) qPCR analysis of relative expression of *Agt* to *Rpl* in tPVAT and (**M**) body weight of flox control and PRDM16cKO mice. (**N**) Mass spectrometry measurements of angiotensinogen levels in serum (n=8 per group) and (**O**) ANGII levels in plasma (n=8 per group) in flox control and PRDM16cKO mice. Individual data points in (K-N) represent data from a single animal, and bars are means ± SEMs. Significance was calculated using unpaired Student’s *t* test. Statistical significance was set at \**P* < 0.05; \*\**P* < 0.01; \*\*\**P* < 0.001; \*\*\*\**P* < 0.0001; ns, not significant.

To determine whether local PRDM16-dependent adipocyte remodeling affects vascular reactivity, we performed pressure myography experiments in mesenteric arteries cleared of PVAT (Fig S2A). Indeed, pressure myograph measurements revealed a substantial increase in ANGII-mediated contraction in PRDM16cKO mice compared to littermate controls (Fig 2A). In contrast, phenylephrine-mediated contraction (Fig 2B), myogenic tone (Fig S2B-D) and endothelial-dependent acetylcholine-mediated relaxation (Fig 2C) were not affected in PRDM16cKO arteries, consistent with an ANGII-specific remodeling of vascular smooth muscle cell (VSMC) reactivity. The increased ANGII-dependent reactivity in PRDM16cKO mice was measured in the absence of PVAT, suggesting persistent remodeling and increased arterial sensitivity, without changes in angiotensin receptor expression (Fig 2D). Furthermore, ablation of *Prdm16* was specific to adipose tissue, with no alteration in arteries (Fig 2E, F). These findings support the existence of an adipocyte-to-VSMC crosstalk that modulates the response to ANGII.

**Fig 2.**
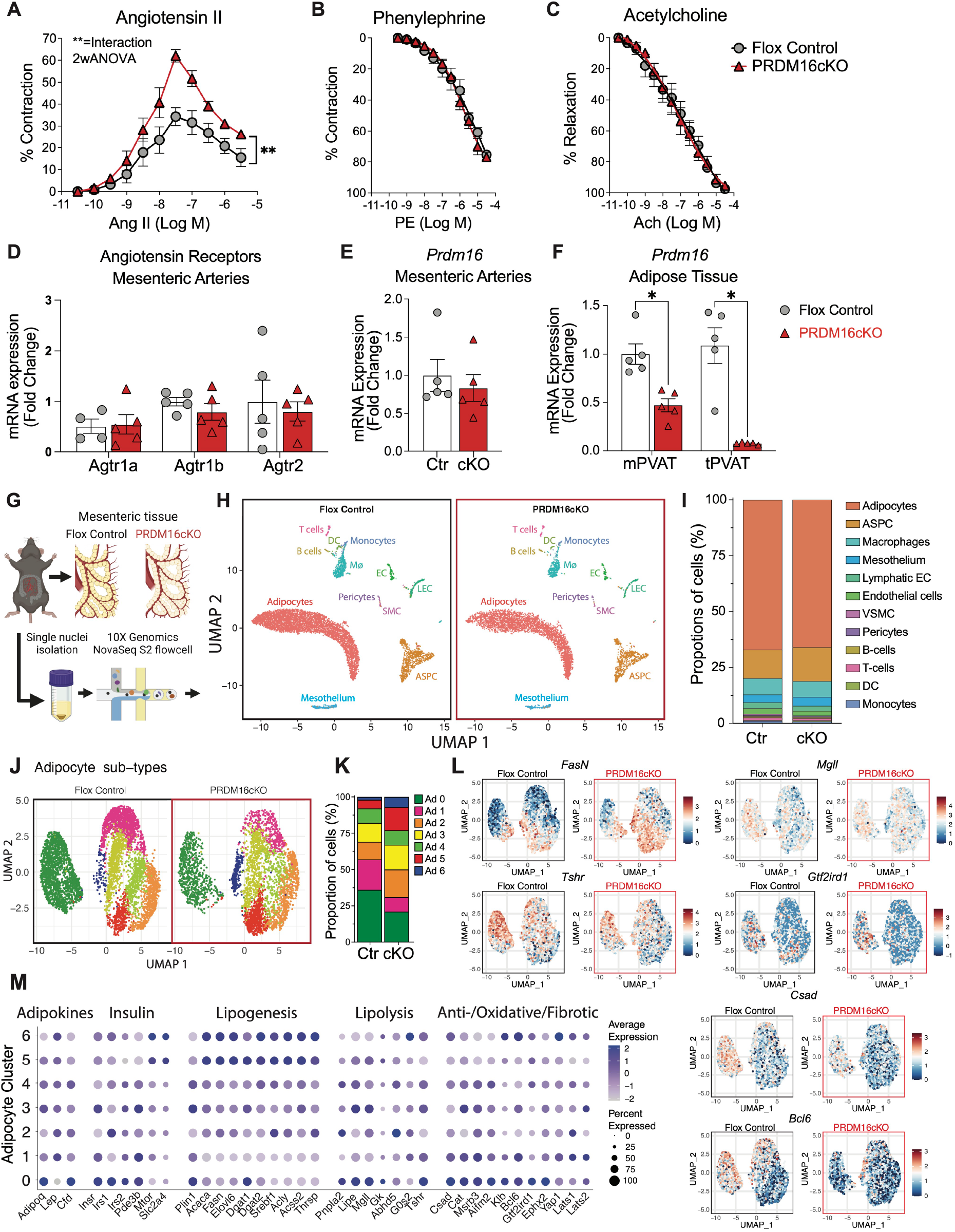
Ablation of beige fat results in increased vasoconstriction of mesenteric arteries to ANGII and remodeling of mesenteric PVAT. **(A-C)** Pressure myography of mesenteric arteries after removal of perivascular fat was conducted in flox control and PRDM16cKO mice (n=4-5 per group; 1-2 mesenteric arteries per mouse) treated with (A) ANGII, (B) phenylephrine and (C) acetylcholine. Data are mean+/-SEM. Statistical analysis was performed using two-way analysis of variance (ANOVA). (**D-F**) Quantitative real-time PCR for (D) *Agtr1a, Agtr1b* and *Agtr2* and(E) *Prdm16* on mesenteric arteries and (F) mPVAT (n=4-5 per group). Individual data points represent data from a single animal, and bars are means ± SEMs. (**G**), Schematic workflow for single nuclei RNA Sequencing of mPVAT. (**H**) UMAP projection of clusters formed by 13,164 murine mPVAT split by genotype. (**I**) Proportion of nuclei in each cluster split by genotype. (**J**) UMAP projection of adipocyte cluster formed by 8783 nuclei from mesenteric adipocytes, split by genotype. (**K**) Proportion of adipocyte nuclei in each adipocyte cluster split by genotype. (**L**) FeatureBlot of different genes split by genotype and (**M**) dot blot of gene expression associated with adipokine secretion, insulin signaling, lipid handling and fibrosis across mesenteric adipocyte subclusters. Statistical significance for all test was set at \**P* < 0.05; \*\**P* < 0.01; \*\*\**P* < 0.001; \*\*\*\**P* < 0.0001; ns, not significant.

### Loss of *Prdm16* results in mesenteric adipocyte remodeling

Mesenteric arteries are resistance arteries critically involved in blood pressure regulation (42,43) and commonly used to assess vascular reactivity. Nevertheless, our understanding of the cellular and molecular landscape of mesenteric PVAT is limited. To further characterize PRDM16-dependent remodeling and its impact on arterial contraction we performed single nucleus RNA sequencing (snucRNASeq) of mesenteric tissue from PRDM16cKO and littermate control mice (Fig 2G). After removing doublets and ambient RNA, a total of 13,164 nuclei were analyzed. Clustering and marker gene analysis identified a total of 15 cell types, including immune cells (macrophages, dendritic cells, monocytes, T- and B-cells), vascular cells (blood and lymph endothelial cells, pericytes and VSMC) and progenitor cells (adipose stem and progenitor cells and mesothelial cells) as well as a large mature adipocyte cluster (Fig 2H, I). Previous studies have highlighted the impact of immune cells on blood pressure regulation (44); however, we did not detect genotype-dependent changes in the immune cell composition of mPVAT by snucRNA Seq (Fig 2H, I), or flow cytometry (Figure S3A-C).

To identify PRDM16-dependent changes in mesenteric adipocytes that could lead to dysregulation of adipocyte-VSMC crosstalk we subclustered mature adipocytes and found that 4 out of 7 detected adipocyte clusters were substantially changed in PRDM16cKO (Fig 2J-M). Besides an increase in adipocytes with a lipogenic signature (*Fasn, Elovl6, Acly* and *Acss2* in Ad5 and Ad6), *Prdm16* deletion also resulted in an approximately 50% reduction in adipocytes associated with lipolysis (*Lipe, Mgll, Tshr* in Ad0 and Ad1), protection from oxidative stress (*Csad, Bcl6, Cat, Msrb3* in Ad0 and Ad1) and a beneficial adipokine profile (*AdipoQ, Cfd* in Ad0 and Ad1) (Fig2 J-M). Furthermore, loss of *Prdm16* affected genes that regulate adipose tissue fibrosis including decreased expression of the anti-fibrotic *Gtf2ird1*, a direct interaction partner of PRDM16 (45), and increased *Yap* (46), previously associated with adipose tissue fibrosis (47) and vascular stiffness (48) (Fig2 J-M), known risk factors for hypertension (49). These findings led us to investigate the impact of PRDM16-dependent adipocyte remodeling on adipose tissue and vascular fibrosis.

### Loss of beige adipocytes leads to vascular remodeling and elevated blood pressure

Picrosirius red (Fig 3A, B) and immunofluorescence (IF) staining for COL1α1 (Fig 3C) revealed increased fibrosis of mesenteric arteries in PRDM16cKO mice. Similarly, aortic and tPVAT RNA (Fig 3D, Fig S4A) and protein analysis (Fig 3E) and COL1α1 IF (Fig 3F) confirmed fibrotic remodeling in the vasculature of PRDM16cKO mice. Similar to mPVAT (Fig 2H, I and Fig S3A-C), we did not detect genotype-dependent changes in immune cell composition in aPVAT (Fig S3D) and only a slight increase in CD206+ macrophages in tPVAT (Fig S3E). In line with our pressure myograph results, these data suggest a crosstalk of adipocytes with VSMC mediating vascular remodeling in PRDM16cKO mice.

**Fig 3.**
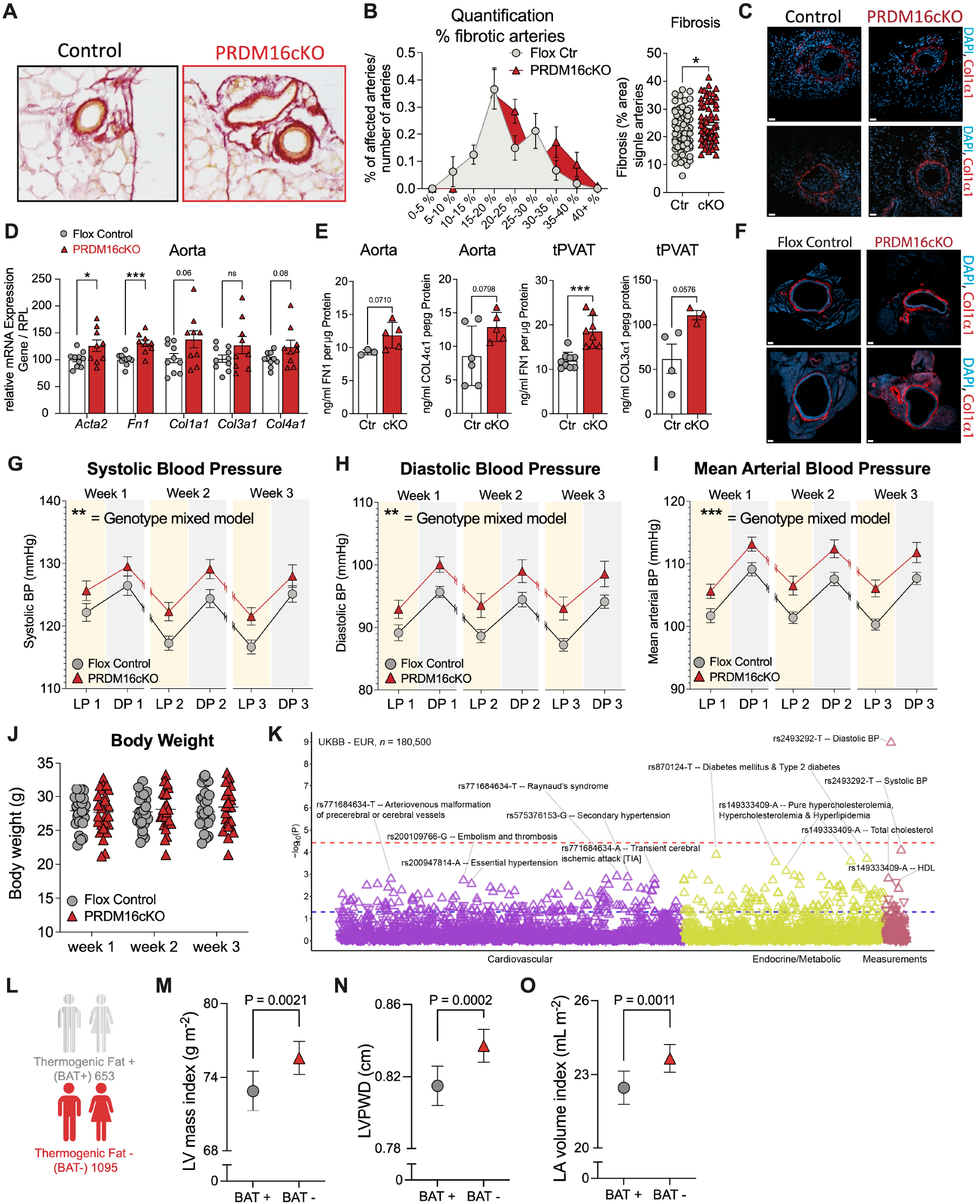
Loss of beige adipocytes leads to vascular fibrosis and elevated blood pressure. (**A**) Representative images of picrosirius red staining on mesenteric arteries and (**B**) quantification of % fibrotic arteries/number of arteries per mouse (n=9 per group) and for total single arteries (right). (**C**) Representative immunofluorescence images for Col1α1 and DAPI nuclei staining on mesenteric arteries. Scale bar 30µm. (**D**) qPCR for *Acta2, Fn1, Col1a1, Col3a1* and *Col4a1* on Aorta of flox control (n=10) and PRDM16cKO (n=9) mice (combined 2 independent experiments). (**E**) ELISA for FN1 (n=3-5) and COL3α1 in Aorta and FN1 (n=8-9 per group) and COL3α1 (n=3-4) in tPVAT of flox control and PRDM16cKO mice. Individual data points in (D, E) represent data from a single animal, and bars are means ± SEMs. Significance was calculated using unpaired Student’s *t* test. **F**) Representative immunofluorescence images for Col1α1 and DAPI nuclei staining on the thoracic aorta. Scale bar 100µm (**G-I**) Measurements of (G) systolic, (H) diastolic and (I) mean arterial blood pressure recorded with implanted radiotelemetry devices in freely moving mice (n=28-29 per group combined from 3 independent experiments). Data are mean+/-SEM. A linear mixed model for repeated measures over time was used to analyze the radiotelemetry data. (**J**) Recording of body weight after transplantation of radiotelemetry devices. (**K**) Association of 25 identified missense variants in exon 9 of *PRDM16* with cardiovascular and endocrine/metabolic traits in the UK BioBank (UKBB). The direction of the triangles indicates the direction of effect (upward: increased risk or level, downward: decreased risk or level). Red and blue dashed lines represent the false discovery rate-adjusted and nominal significance *P* value thresholds, respectively. (**L-O**) Echocardiogram measurements from a cohort of 653 patients with (BAT+) and 1095 patients without (BAT-) thermogenic adipose tissue derived from a previously described propensity-matched cohort (15). (L) Left ventricular (LV) mass index (LV mass/body surface area), (M) left ventricle posterior wall diameter and (N) left atrium (LA) volume index (LA volume/body surface area). Error bars indicate 95% CI. Data was assessed using a general linear model (GLM), and least square means (LSMeans) were determined adjusting for age, sex, body-mass-index, race, and outdoor temperature. Pairwise comparisons were performed using the least significant difference (LSD) test, and 95% confidence intervals were reported. Analyses were conducted using SAS software (SAS Institute, Cary, NC, USA). Statistical significance for all test was set at \**P* < 0.05; \*\**P* < 0.01; \*\*\**P* < 0.001; \*\*\*\**P* < 0.0001; ns, not significant.

Both increased arterial ANGII-reactivity and vascular remodeling influence blood pressure regulation, leading us to investigate systemic blood pressure in PRDM16cKO mice. We implanted radiotelemetric monitoring devices in PRDM16cKO and littermate control mice and recorded blood pressure in freely moving animals over a three-week period (Fig 3G-I and Fig S4B). Strikingly, PRDM16cKO mice showed a significant elevation of systolic (Fig 3G), diastolic (Fig 3H) and mean arterial blood pressure (Fig 3I) both during the light and active dark phases. This highlights the role of PRDM16-dependent adipocyte remodeling on blood pressure regulation, independent of body weight (Fig 3J) or any other detectable metabolic abnormality.

Increased myosin light chain phosphorylation in mesenteric arteries of PRDM16cKO mice, a surrogate for arterial contraction, further supported these findings (Fig S4C, D). Although heart rate was slightly increased in PRDM16cKO mice, particularly during the first week of recording (Fig S4B), cardiac function assessed by echocardiography did not demonstrate any changes (Fig S5A, B), nor did we detect signs of cardiac fibrosis by histology (Fig S5C). Similarly, we did not detect any overt kidney pathology (Fig S5D, E). Adjusting blood pressure readings for the observed changes in heart rate, confirmed a heart rate-independent, persistent increase in blood pressure in PRDM16cKO mice (Extended Table 1). Taken together these results indicate that PRDM16-dependent ablation of beige adipocyte identity, independent of body weight or insulin resistance, leads to a significant increase in blood pressure.

We previously showed that humans with detectable brown fat have significantly decreased odds of hypertension, compared to matched controls lacking brown fat (15). Moreover, a meta-analysis identified a coding variant in exon 9 of *PRDM16* (rs2493292), the same exon deleted in PRDM16cKO mice, to be associated with hypertension in humans (50). We confirmed the association of rs2493292-T in *PRDM16* with increased diastolic and systolic blood pressure in the UK Biobank (UKBB, 180,500 individuals) (Fig 3O) and in patients with European Ancestry (EUR) in the Mount Sinai Million Health Discoveries Program (MSM-EUR, 21,680 individuals), which also revealed an increased association with thoracic aneurysm (Fig S4E). Further, we identified rs371654192-C in the MSM-EUR to be associated with increased systolic blood pressure and morbid obesity (Fig S4E), aligning well with findings from PRDM16cKO mice (Fig 4G) (23). Additionally, rs200947814-A and rs575376153-G were associated with an increased risk for essential and secondary hypertension in UKBB, respectively (Fig 3O). Interestingly, analysis of patients with African Ancestry in MSM (MSM-AFR, 19,856 individuals) showed a positive association between *PRDM16* variants and pulmonary hypertension (rs199998420-A and rs114204766-A). We also identified a *PRDM16* variant positively associated with combined systolic and diastolic heart failure (rs201807364-C) in MSM-AFR, which was also associated with reduced systolic blood pressure (Fig S4F).

**Fig 4.**
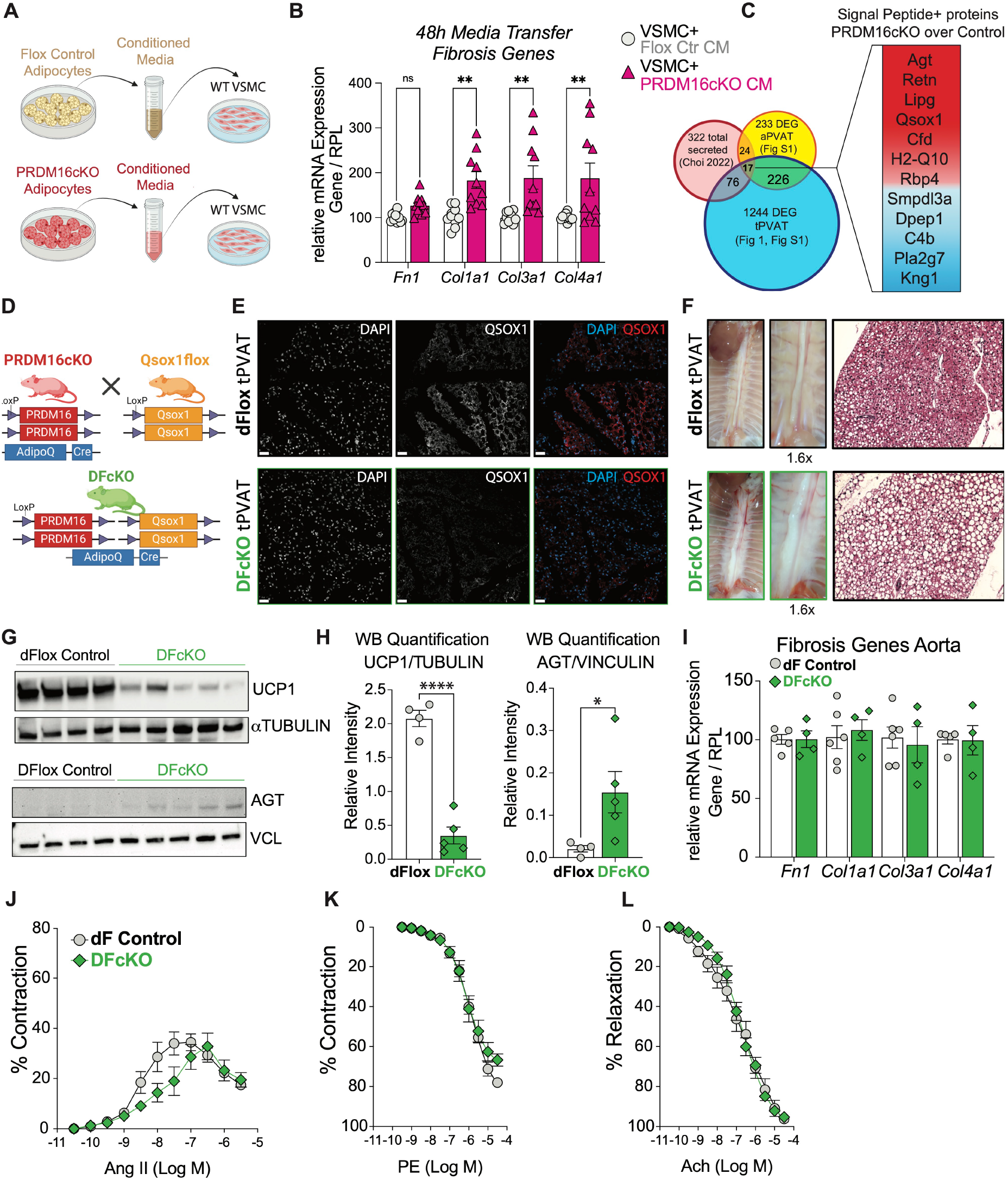
Loss of QSOX1 in PRDM16cKO adipocytes rescues aortic fibrosis and ANGII-mediated hypercontraction. (**A**) Schematic of media shuffling experiment setup. (**B**) qPCR for *Fn1, Col1a1, Col3a1* and Col4a1 on wild type primary aortic vascular smooth muscle cells (VSMC) treated for 48h with conditioned media from flox control or PRDM16cKO adipocytes. Individual data points represent data from a single well from 3 independent experiment, and bars are means ± SEMs. (**C**) Venn diagram of DEGs from tPVAT and aPVAT of PRDM16cKO mice overlayed with adipocyte-derived secreted proteins identified previously(54). 12 genes are identified to be down (blue) or upregulated (red) in PRDM16cKO mice and signal peptide P-positive. (**D**) Schematic of generation of double flox adipocyte conditional knock-out (DFcKO) mice (**E**) Immunofluorescence image for QSOX1 and nuclei staining DAPI in tPVAT of double PRDM16flox; Qsox1flox control (dF control) and DFcKO. Scale bar 30µm. (**F**) Appearance and HE staining of the thoracic PVAT in dFlox control and DFcKO mice. (**G, H**) Representative western blot of tPVAT for UCP1 (top), αTUBULIN (top) and AGT (bottom) and VINCULIN (bottom) and (H) quantification. Statistical analysis was performed using *Student’s* T-Test. (**I**) qPCR for *Fn1, Col1a1, Col3a1* and *Col4a1* on the thoracic aorta of dFlox Control and DFcKO mice. Individual data points represent data from a single animal, and bars are means ± SEMs. (**J-O**) Pressure myography of mesenteric arteries after removal of perivascular fat was conducted in dflox control and DFcKO mice (n=5 per group; 1-2 mesenteric arteries per mouse) treated with (J) ANGII, (K) phenylephrine and (L) acetylcholine. Data are mean+/-SEM. Statistical significance for all test was set at \**P* < 0.05; \*\**P* < 0.01; \*\*\**P* < 0.001; \*\*\*\**P* < 0.0001; ns, not significant.

As a direct clinical readout of long-term chronic elevated blood pressure-induced end-organ damage (6,51–53), we reviewed echocardiograms from our human brown fat cohort and assessed left ventricular (LV) mass index (LV mass/body surface area), left ventricle posterior wall diameter, and left atrium volume index (LA volume/body surface area) (Fig 3L-O). In a cohort of 653 patients with (BAT+) and 1,095 without brown fat (BAT-) (Fig 3L), we found that LV mass index (Fig 3M), LV posterior wall diameter (Fig 3N) and left atrium volume index (Fig 3O) were significantly increased in BAT-patients. Together with the elevated blood pressure in PRDM16cKO mice, these data highlight a previously unknown and physiologically important crosstalk between thermogenic adipocytes and the vasculature, presumably VSMC, regulating blood pressure in humans and mice.

### An adipocyte-derived secreted factor is sufficient to modulate vascular remodeling

To identify how *Prdm16*-deficient adipocytes directly induce vascular remodeling and whether this is through a secreted factor, we established a cell culture conditioned media transfer system from primary adipocytes to primary aortic VSMC (Fig 4A). Treatment of wild-type primary aortic VSMC with conditioned media from PRDM16cKO adipocytes, in comparison to floxed control adipocytes, was sufficient to significantly induce ECM gene expression in VSMC (Fig 4A, B). By overlaying a previously reported dataset of proteins secreted by primary adipocytes (54) with differentially expressed genes (DEGs) from PRDM16cKO PVAT RNA sequencing (Fig 1G, S1E-H), we identified 17 genes present in all three datasets. Analysis for signal peptide-positive genes, as a measure of actively secreted proteins, resulted in a list of 12 PRDM16-regulated genes that are predicted to be secreted from adipocytes. As expected from our RNA sequencing, *Agt* was one of the PRDM16-regulated secreted factors. In addition to other genes known for their metabolic role, such as *Retn* and *Cfd* (Fig 4c), we identified *Quiescin sulfhydryl oxidase 1* (*Qsox1*) as an adipocyte-derived, PRDM16-regulated secreted enzyme (Fig 4C, Fig S6A-C). Analysis of published PRDM16 ChIP sequencing data (39) showed binding of PRDM16 close to the transcriptional start site and in intronic regions of *Qsox1*, with PRDM16cKO showing a substantial increase in H3K4me1 in the second intron of *Qsox1* and a modest increase in H3K4me2 (Fig S6A). The substantial increase in H3K4me1 suggests an intronic enhancer that is repressed by PRDM16, and which may become de-repressed in PRDM16cKO adipocytes (Fig S6A). Consistent with these epigenetic data, analysis of *Qsox1* expression in different adipose depots showed that adipocyte-specific deletion of *Prdm16* resulted in a significant upregulation of *Qsox1*, primarily in thermogenic adipose tissue (Fig S6B). Similarly, loss of thermogenic adipocyte function in obesity led to increased *Qsox1* expression (Fig S6C). QSOX1 catalyzes disulfide bridge formation in the extracellular space and regulates ECM assembly in cancer (55). Strikingly, single nucleotide polymorphisms in human *QSOX1* are associated with blood pressure (56), and whole-body *Qsox1* knock-out mice have lower blood pressure (57). The role of QSOX1 in adipocytes has not been studied before and the impact of adipocyte-derived QSOX1 on vascular physiology remains completely unknown.

### *Qsox1* is required to provoke vascular dysfunction in PRDM16cKO mice

To investigate the impact of QSOX1 on PRDM16-mediated PVAT remodeling and vascular function we generated *Qsox1* floxed mice (Fig S6D) and crossed them to *AdipoQ*-Cre mice for an adipocyte-specific *Qsox1* deletion (Qsox1cKO) (Fig S6D, E). Adipocyte-specific loss of *Qsox1* alone had no impact on body or adipose tissue weight, insulin resistance or cold tolerance on a standard diet (Fig S6F-L). We then generated mice with an adipocyte-specific double deletion of *Prdm16* and *Qsox1* (double floxed, *AdipoQ*-Cre+ mice -DFcKO) to address whether QSOX1 is required for vascular fibrosis and increased ANGII-mediated vasoconstriction in PRDM16cKO mice. IF staining showed a clear loss of QSOX1 in adipocytes of DFcKO compared to double floxed control (dFlox) mice (Fig 4E). Similar to PRDM16cKO mice, DFcKO mice exhibited ‘whitened’ thoracic (Fig 4F) and abdominal PVAT (Fig S7A) with enlarged adipocytes and reduced UCP1 expression (Fig 4G, H, Fig S7B), without effects on body weight (Fig S7C). In addition to a lack of UCP1, DFcKO still demonstrated increased *Agt* RNA (Fig S7D) and protein expression (Fig 4G, H) and increased ALDH3B2 and REEP6 (Fig S7E, F), two white adipocyte marker genes, as compared to dFlox control mice. Taken together, deletion of *Qsox1* in addition to *Prdm16* does not rescue the PRDM16cKO-specific loss of thermogenic signature in DFcKO mice, nor does it correct the elevated local tissue AGT protein levels (Fig 4G, H) seen in single PRDM16cKO mice (Fig 1J). Nevertheless, in contrast to PRDM16cKO mice, DFcKO mice showed no differences in ECM gene expression in the aorta compared to dFlox littermate controls (Fig 4I). Encouraged by this result, we performed pressure myography on DFcKO and dFlox control mesenteric vessels and strikingly found a complete rescue of ANGII-mediated arterial contraction in DFcKO mice (Fig 4J), as compared to the ANGII-mediated hypersensitivity in single PRDM16cKO arteries (Fig 2A). Similar to PRDM16cKO mice, we did not detect any changes in phenylephrine-mediated contraction (Fig 4K), acetylcholine-mediated relaxation (Fig 4L), or myogenic tone (Fig S7G-I) between DFcKO and dFlox control mice.

Taken together, we discovered a novel PRDM16-dependent adipocyte-to-VSMC crosstalk regulating ANGII-dependent vascular reactivity, fibrosis and blood pressure. We identify QSOX1 as an adipocyte-derived secreted protein, that is dramatically upregulated in PRDM16cKO mice, and simultaneous deletion of *Qsox1* and *Prdm16* rescues vascular reactivity and remodeling observed in PRDM16cKO mice. Overall, our findings describe a new physiologically important crosstalk between adipocytes and the vasculature that is independent of obesity, insulin resistance or thermogenic activity, highlighting the contribution of adipocyte identity on cardiovascular outcomes.

## DISCUSSION

White adipose tissue accumulation in obesity and aging is associated with increased blood pressure, however, the impact of adipocyte identity, independent of systemic insulin resistance and inflammation has been unclear. Clinical studies showed that individuals lacking active brown fat have higher odds of hypertension independent of body weight (15), suggesting a beneficial impact of thermogenic brown adipose tissue. In humans, even mild chronically elevated blood pressure is associated with end organ damage (3–7), and indeed we found that humans lacking detectable thermogenic brown fat showed remodeling of the left ventricle and atrium, clinical sequelae of long-term blood pressure elevation (6,51–53) and independent predictors of hypertension (53,58). Adult human thermogenic fat shares more similarities with rodent beige fat than classical brown fat (20–22). Therefore, mice with a PRDM16-dependent ablation of beige fat represent a useful model to study the impact of human thermogenic adipose tissue on vascular physiology.

Indeed, we and others identified coding variants in exon 9 of *PRDM16* to be associated with hypertension and its comorbidities (Fig 3K, Fig S4E,F) (50). Our mouse data, support these findings and demonstrate that adipocyte-specific loss of *Prdm16* (23) is sufficient to induce elevated blood pressure by modulating ANGII-mediated arterial contraction in mice. In contrast, VSMC-specific deletion of PRDM16 results in reduced blood pressure and vascular reactivity (59), highlighting an important and adipocyte-specific effect of PRDM16 on blood pressure regulation, consistent with the observed increased risk for hypertension in humans (50). In mice ablation of beige adipocytes is also associated with an impaired repair response to endovascular injury (14). Further, QSOX1 is increased in the intima after injury, and treatment of VSMC with QSOX1 induced migration (60) supporting a potential role for extracellular QSOX1, such as that derived from adipocytes on VSMC physiology. Consequently, knockdown of *Qsox1* in endovascular injury resulted in reduced neointima formation (60), together suggesting a more general role for adipocyte-derived, PRDM16-regulated QSOX1 in adipocyte-to-VSMC crosstalk.

As a consequence of beige fat deletion, PRDM16cKO mice demonstrate reduced thermogenesis and increased ANGII reactivity. Previous work showed that thermogenic fat can blunt noradrenaline-induced arterial contraction in control (12) but not ANGII-treated mice (61). However, increased thermogenesis after β3-adrenergic activation was able to reinstall anticontractile effects in ANGII-treated rats involving the induction of anti-oxidative genes *Cat* and *Sod2* (61). Genetically increased thermogenesis, however, did not affect blood pressure (34), suggesting a role of beige adipocytes in vascular reactivity beyond thermogenesis. In line with this, DFcKO mice showing a similar reduction in thermogenesis as PRDM16cKO mice had normalized ANGII-mediated contraction.

Remodeling of PVAT in PRDM16cKO and DFcKO mice was accompanied by increased tissue levels of AGT and adipocyte-specific AGT signaling has been associated with obesity-mediated hypertension (40,41), while adipocyte-specific deletion of *Agt* prevents obesity-induced hypertension (62). Genetic overexpression of *Agt* increases blood pressure but also circulating levels of ANGII (40), which was not observed in our PRDM16cKO mice. Instead, we observed a local increase in AGT without effects on circulating AGT or ANGII levels, suggesting a “slow pressor” effect of ANGII (63). This model posits that small sub-pressor doses of continuous ANGII can raise blood pressure without a concomitant increase in plasma levels, potentially through increased oxidative stress (63). Both ANGII- and obesity-associated hypertension are associated with increased oxidative stress and vascular stiffness (34,64,65), and similar to our adipocyte-specific snucRNA seq showing loss of anti-oxidative gene expression (*Csad, Bcl6, Cat, Msrb3*), deletion of *Prdm16* in neural stem cells and cardiomyocytes resulted in increased oxidative stress and ROS production (66,67). Loss of antioxidative enzymes in PVAT of diabetic rats was shown to affect vasoreactivity (68), and VSMC-specific overexpression of *Bcl6* attenuates vascular remodeling and blood pressure in hypertensive rats (69). QSOX1, through suppression of anti-oxidative mediators, has been shown to increase intracellular ROS levels in cancer cells (70), perhaps potentiating oxidative stress induction by *Prdm16* ablation. Ablation of *Qsox1* in addition to *Prdm16* in DFcKO mice may counteract the AGT-mediated effect through regulation of oxidative stress.

Whole body deletion of *Qsox1* alone results in lower blood pressure (57), while our adipocyte-specific deletion of *Prdm16* led to a dramatic increase in *Qsox1* and increased ANGII-dependent vascular reactivity and blood pressure. Simultaneous deletion of *Qsox1* and *Prdm16* rescued the increased vascular fibrosis and ANGII-dependent increased reactivity observed in single *Prdm16* knockout mice. QSOX1 catalyzes the formation of intra- and extracellular disulfide bonds (55,71) and oxidative post-translational modification of cysteine thiols in proteins can regulate their structure and function (72). Strikingly, oxidative reduction of disulfides in the angiotensin receptor by DTT treatment results in ablation of ANGII-mediated contraction in the rabbit aorta (73) supporting the hypothesis that increased QSOX1 observed in PRDM16cKO mice, through direct disulfide bond formation, could affect angiotensin receptor sensitivity on VSMC. Finally, QSOX1 can directly regulate ECM composition and cell migration in cancer (55,74) and QSOX1 plasma levels positively correlate with fibrosis in metabolic dysfunction-associated steatotic liver disease (75) aligning well with our finding of increased vascular fibrosis in PRDM16cKO mice.

Taken together, this study highlights the importance of adipocyte identity and the potential of targeting QSOX1 to prevent vascular fibrosis and elevated blood pressure in lean individuals with increased visceral adiposity, who may not have elevated circulating ANGII levels. Inhibition of QSOX1 could represent a novel strategy to prevent vascular stiffness and lower blood pressure, while activation of beige adipocytes could provide previously unappreciated benefits for cardiovascular health.

## Acknowledgments

We thank Xiaojing Huang and Yue Liu for experimental advice and discussions. We thank Hong Duan and other members from The Rockefeller Genomics core and Matt Paul, Thomas Carroll, Ji-Dung Luo and other members from The Rockefeller Bioinformatics Resource Center for performing, analysis and visualization of snucRNA Seq and RNA Seq data. We thank Caroline S Jiang and Roger Vaughn from the Rockefeller Center for Clinical and Translational Science for support in statistical analysis. This work was further supported by the Flow Cytometry Resource Center, the Bio-Imaging Resource Center, the CRISPR and Genome Editing Center and the Transgenic and Reproductive Technology Center and the Laboratory of Comparative Pathology at the Rockefeller University. We thank all members of the Laboratory for Molecular Metabolism and members of our Leducq funded Network for lively discussions and intellectual exchange. Figures 2G, 3K, 4A, D and S6D were created with Biorender (https://biorender.com). *Created in BioRender. Koenen, M*. (2025) https://BioRender.com/exd5yys. This research was conducted using the UK Biobank resource under application number 53074.

## Funding

The Alexandrine and Alexander L. Sinsheimer Foundation (PC)

National Institute of Diabetes and Digestive and Kidney Diseases (RC2DK129961) (PC) National Institute of Health (R01HL 152195) (ADL)

Leducq Foundation (21CVD01) (PC, YI, MEK, ACM and RMT) Women&Science Initiative at Rockefeller University (MK)

German Research Foundation (Walter Benjamin KO 6368/1-1) (MK) Charles H. Revson Foundation (Grant No. 23-22) (MK)

Kellen Women’s Entrepreneurship Fund with support from Lulu Wang (MK).

National Center for Advancing Translational Sciences, NIH, through The Rockefeller University (Grant UL1TR001866) (TB and NGB).

National Center for Advancing Translational Sciences, Clinical and Translational Science Awards (CTSA) grant UL1TR004419. This work was supported in part through the computational and data resources and staff expertise provided by Scientific Computing and Data at the Icahn School of Medicine at Mount Sinai.

## Author contributions

PC conceptualized the study with contribution from MK and TB. PC and MK acquired funding, MK visualized data, MK, TB designed and interpreted experiments MK, TB, GP, LR, ACM, ZL, RA, IDG, SJH, ZFHC and SDB performed experiments. JRO performed snucRNASeq analysis and visualization. JAB, NGB and MEK analyzed human data. XZ analyzed and visualized ChiP Seq data. PC, MK, ADL, RMT, YI, SDB, NGB supervised study/study segments, PC, SDB, IDG, ADL, ACM, RMT, MEK and YI provided technical and intellectual input. MK and PC wrote the manuscript, and all authors edited it.

## Competing interests

TB is an employee and shareholder of Roche Diagnostics International, Rotkreuz, Switzerland.

All other authors declare that they have no competing interests.

## Supplementary Materials

Materials and Methods Figs. S1 to S7

Tables S1 to S2

