## Supplement_Materials_Methods_Figures for "Ablation of *Prdm16* and beige fat causes vascular remodeling and elevated blood pressure"

**The PDF file includes:**

Materials and Methods
Figs. S1 to S7
Tables S1 to S2

### Materials and Methods

Mouse Models

Prdm16cKO mice were generated as previously described (23) by crossing *AdipoQ-Cre* mice (JAX 028020) with *Prdm16*$^{loxP/loxP}$ mice (JAX 024992). Qsox1$^{loxP/loxP}$ mice were generated at the CRISPR/Genome Editing Resource Center (CGERC) and the Transgenic/Reproductive Technology Resource Center (TRTRC) of the Rockefeller University, New York, NY, USA. To generate adipocyte-specific Qsox1cKO we crossed *Qsox1*$^{loxP/loxP}$ mice with *AdipoQ-Cre* mice (JAX 028020). To generate DFcKO mice with an adipocyte-specific deletion of *Prdm16* and *Qsox1*, we crossed PRDM16cKO and *Qsox1*$^{loxP/loxP}$ mice. All animals in this study were male mice on a C57BL/6J background and maintained on a 12h light/dark cycle with free access to food and water. If not indicated differentially, all mice were between 11-14 weeks old, housed at 23°C and fed with a standard rodent chow diet. For cold exposure experiments, mice were placed at 8°C or for thermoneutrality at 30˚C for 1 week with one mouse per cage. Animal care and experimentation were performed according to procedures approved by the Institutional Animal Care and Use Committee at the Rockefeller University. Mice were randomly assigned cages, independent of genotype and given a new experimental number for each experiment to ensure blinding of genotype. Exception for representative western blots, after blinded protein isolation and quantification, mice were organized in groups dependent on genotype on the SDS page.

Blood Pressure Measurements

Blood pressure was measured by implantable radio transmitters as previously described (76). Briefly, male mice at age 11-14 weeks were housed in pairs. Prior to implantation, the animals received buprenorphine (0.8 mg/Kg, s.c.) and a transmitter was implanted (model HD-X11, Data Sciences International, St. Paul, MN, USA), while mice were under isoflurane anesthesia. The catheter tip was positioned in the thoracic aorta via the left common carotid artery and the body of the probe was inserted subcutaneously in the dorsal right flank. Animals were allowed to recover for 5–7 days. Heart rate (HR), systolic blood pressure (SBP), diastolic blood pressure (DBP), and mean arterial pressure (MAP) were collected weekly over a 24h period in freely moving conscious mice in their cages. Pressure traces were checked and animals with dampening of the haemodynamic profile were excluded from the analysis.

Analysis of Blood Pressure Readings

For each mouse, systolic, diastolic and mean arterial blood pressure as well as heart rate was recorded. The mean for each parameter was calculated for dark and light cycles each for a 12h period. Data from 3 experiments were pooled and used for the final analysis. A linear mixed model for repeated measures over time (SAS Proc Mixed) was used to analyze the radiotelemetry data with fixed effects of genotype (PRDM16cKO vs. Flox Control), week treated as a categorical factor, and the interaction between genotype and week. This method prevented list-wise deletion due to missing data. Unstructured covariance structure was chosen with the lowest corrected Akaike's information criteria and Bayesian information criteria. Heart rate was included as a covariate for the blood pressure models and the adjusted mean difference between PRDM16cKO vs. Flox Control mice with 95% confidence intervals were reported.

Pressure Myograph

Mesenteric arteries were collected from 12-13-week-old male mice. Third order mesenteric arteries (MA) were cleaned of surrounding fat and vascular reactivity experiments were performed as described previously (77). Briefly, mesenteric arteries (MA) were mounted on glass cannulas in a pressure myograph chamber (Danish MyoTechnology, Aarhus, Denmark). The vessel orientation in relation to the flow *in vivo* was maintained. Vessel viability was maintained using Krebs solution (in mM: NaCl 118, KCl 4.7, $MgCl_2$ 1.2, $KH_2PO_4$ 1.2, $CaCl_2$ 2.5, $NaHCO_3$ 25 and glucose 10.1), at 37 °C and oxygenated (95% $O_2$ and 5% $CO_2$). A pressure interface controlled intraluminal pressure and flow in MA. The vessel diameter was monitored in real time using a microscope connected to a digital video camera (IC Capture) and computer software VediView 1.2 (Danish MyoTechnology, Aarhus, Denmark) with edge detection capability. MA were equilibrated for 15 min at 80 mm Hg, pre-constricted with PE (1 µM) and a cumulative concentration-response curve of Ach (0.1 nM – 30 µM) was performed to evaluate the integrity and the function of the endothelium. MA with less than 80% vasodilatory response to Ach were discarded. Vascular smooth muscle functions were evaluated by performing cumulative concentration-response curves of PE (1 nM – 30 µM), ANGII (0.1 nM – 1 µM) and myogenic tone.

<u>Mass Spectrometry</u>

12 µl of serum sample were incubated with 12 µL of 20mM dithiothreitol (DTT) and 16 M urea (final 8M urea, 10 mM DTT, 50mM ammonium bicarbonate (ABC)) for 1h at room temperature (RT). Another 6 µL of 50mM DTT were added to each of the samples (in 50 mM ABC, no urea) to ensure complete reduction. The samples were then alkylated by addition of 20 µL of 75mM iodoacetamide (IAA; 30mM final in ABC) in the dark at RT for 1.5h. 6 µg of LysC was added to each of the sample. The samples were then allowed to digest at 28° C overnight with shaking. The following day, the samples were further diluted with 60 µL of 50mM ABC. 10 µg of trypsin 372 was added to each of the samples and samples were digested with trypsin for 6 hours at 28° C 373 at 1400 rpm. The samples were then quenched with 10 µL of 10% TFA (pH approximately 1). 5 µL of sample was loaded onto 4 C18 empore discs. The cleaned-up samples were speed vacuumed and re-dissolved in 20 µL of 0.1% LC-grade formic acid. 1 uL was loaded onto a reverse-phase nano-LC-MS/MS (EasyLC 1200, Fusion Lumos, Thermo Fisher Scientific) for analysis. Peptides were separated with a gradient in which the proportion of buffer B (H2O with 0.1% formic acid) in buffer A (acetonitrile with 0.1% formic acid) increased from 2% to 90% over 79 minutes (300 nl/min flow rate). MS and MS/MS data were recorded at resolutions of 60.000 and 30.000 with AGC's of 1,000,000 and 50,000 respectively. Parallel reaction monitoring (PRM) was used to target peptides unique to AGT (AIQGLLVTQGGSSSQTPLLQSIVVGLFTAPGFR and LPTLLGAEANLNNIGDTNPR). PRM experiment data were analyzed with SkyLine v.4.2. Samples were analyzed in the following order: KO, WT and values were normalized to WT to derive fold-change between the two genotypes.

<u>Echocardiography</u>

Mice were anaesthetized with isoflurane under continuous monitoring and placed on a heating pad of a recording stage connected to a Vevo 2100 ultrasound machine. The anterior chest wall was shaved, ultrasound gel was applied, and electrodes were connected to each limb to simultaneously record an electrocardiogram. Two-dimensional (short axis-guided) M-mode and B-mode measurements (at the level of the papillary muscles) were taken using an 18–32 MHz MS400. For analysis, at least three measurements were averaged; measurements within the same heart rate interval (450 ± 50 bpm) were used for analysis.

Metabolic characterization of mice

For fasting blood glucose measurements and glucose tolerance test (GTT), mice were single housed in the morning in cages with fresh sani-chip bedding and ad libitum access to water but without food. Mice were kept in a procedure room free of noise or vibration throughout the experiment. After 4h, blood was collected from the tail vein, and glucose levels were measured using a glucose meter (Nova Max Plus). Intraperitoneal GTT was performed following a 6h fasting procedure and started by administering at time 0 indicated doses of glucose by intraperitoneal injection. Following injection, blood glucose measurements were taken from the tail vein at indicated timepoints. For cold tolerance test (CTT) mice were single housed in cages with ad libitum food and water and 2g of enrichment per cage. Mice where pre-acclimated for 5 days in thermoneutrality to deactivate any residual thermogenesis at room temperature and transferred to 8˚C. Rectal temperature was measured at timepoint 0 and then every 30 minutes for 5h, followed by measurements after 24h, 48h, 72h and 96h and at the end of the experiment after 7 days of cold exposure. For studies involving diet-induced obesity, mouse body weights were monitored once a week, providing fresh 60% high fat diet (Research Diets) at least once a week.

RNA Sequencing

Mice were sacrificed with $CO_2$ and perfused with 1xPBS to remove blood. tPVAT and aPVAT were collected, snap frozen in liquid nitrogen and stored at -80˚C. Frozen samples were pulverization in a liquid nitrogen pre-cooled metal cell crusher and total RNA was extracted by TRIzol (Invitrogen). Tissue powder was mixed with TRIZol and incubated under constant rotation for 15min at 4˚C for full lysis and centrifuged to pellet cell debris. Supernatant was used to purify total RNA using chloroform, ethanol and RNeasy Micro Kit (Qiagen). cDNA was synthesized from 1 µg of RNA using the High-Capacity cDNA Reverse Transcription Kit (Applied Biosciences). At the Rockefeller Genomics Resource Center RNA quality was measured by the Agilent 2100 bioanalyzer chip and samples prepared with Illumina Stranded mRNA Prep, Ligation (96 Samples) /Catalog 20040534 with IDT for Illumina RNA UD Indexes Set B, Ligation (96 Indexes, 96samples). Sequencing was performed with NextSeq2000 P3: 2x50 @550pM with 12%phix.

RNA Sequencing Analysis

Sequence and transcript coordinates for mouse mm10 genome and ensembl gene models were retrieved from UCSC (GRC Mouse Build 38). Transcript expressions was calculated using the Salmon quantification software (78) (version 1.8.1) and gene expression levels as TPMs and counts retrieved using Tximport (79) (version 1.30.0). Normalization and rlog transformation of raw read counts in genes were performed using DESeq2 (80) (version 1.42.1). For visualisation in genome browsers, RNA-seq reads are aligned to the genome using Rsubread's subjunc method (version 2.8.2) (81) and exported as bigWigs normalised to reads per million using the rtracklayer package (version 1.54.0). GSEA analysis of gene set enrichment was performed using the fgsea (version 1.28.0) with the MsigDB pathway gene sets (version v2022.1.Mm).

Single nuclei RNA Sequencing

Mice were perfused with 1x PBS and after removal of lymph nodes, mesenteric adipose tissue was snap frozen and stored at -80˚C. For single nucleus isolation tissue was homogenized in 3ml TST buffer (146 mM NaCl, 21 mM $MgCl_2$, 10 mM Tris-HCl pH7.5, 1 mM $CaCl_2$, 1% Molecular Grade BSA, 0.2 U/µL RNase inhibitor, 0.1% Tween-20, in Ultrapure $H_2O$) using GentleMACs C-tubes

(Miltenyi 130-093-237). Tissues were homogenized immediately using a gentleMACS Tissue Dissociator (Miltenyi Biotec) by running the "mr_adipose_01" program twice. After homogenization, samples were incubated on ice for 5 min and filtered through a 30 µm MACS SmartStrainer (Miltenyi Biotec 130-110-915) into a new 15 mL falcon tube. The gentleMACS C tube was then rinsed with 3 mL of ST buffer (146 mM NaCl, 21 mM $MgCl_2$, 10 mM Tris-HCl pH7.5, 1 mM $CaCl_2$, 0.2 U/µL RNase inhibitor (Sigma-Aldrich 3335402001), in Ultrapure $H_2O$) and the wash buffer was filtered through the same 70 µm MACS SmartStrainer and combined with the homogenate. Samples were centrifuged at 500g for 5 min in 4˚C and brake set to 5. The pellet was resuspended in 400µl 1xRSBTwS Buffer (Made from 6X RSBTwS Buffer: 60mM NaCl, 18mM $MgCl_2$, 60mM Tris-HCl pH 7.5, 0.6% (w/v) Tween20, 0.2 U/µL RNase inhibitor in ultrapure $H_2O$) and mixed with equal amount of 50% iodixanol (1xRSBTwS buffer with 50% Iodixanol solution) and an iodixanol gradient was formed (25%, 30%, 35%). After centrifugation at 10,000g for 20 min at 4˚C the nuclei pellet was collected at the 30%-35% interphase. The pellet (~50µl) was collected into 1ml of 1xPBS+1%BSA and mix well and filtered through a 10µm filter, wash with 1ml 1xPBS+1%BSA and centrifuged at 300g for 8 min with brake at 5 at 4˚C. The purified nuclei pellet was recovered in 60 µL of resuspension buffer. Nuclei from the same genotype were combined as one sample (n=8 flox control, n=10 PRDM16cKO). The sample libraries were prepared using 10x Chromium Next GEM Single Cell 3' kit v3.1 16 rxns (PN-1000268). The libraries QC were done on TapeStation. Then the 2 libraries were run on Novaseq S2 flowcell (pooled with 4 other libraries, total of 4.1 billion reads) in a 2 x 50 bp paired end sequencing format @ 2.6nM with1%phix at the Rockefeller University Genomics Resource Center.

<u>snucRNAseq Data Analysis</u>

Libraries were aligned to the *Mus musculus* mm10 genome (v2020) using Cell Ranger v6.1.2. Ambient RNA contamination was removed using the DecontX R package (v1.0.0) (82). To identify and exclude doublets, Scrublet v0.2.3 (83) was run in Python and imported into R via reticulate v1.35.0 (84) (reticulate: Interface to 'Python'. R package version 1.42.0, https://github.com/rstudio/reticulate, https://rstudio.github.io/reticulate/.). All downstream analyses were performed with Seurat v5.1.0 (85). Nuclei with >0.5 mitochondrial content were excluded from further analyses. For batch correction, Flox Control and PRDM16cKO nuclei were subset into separate objects and integrated using Seurat's anchor-based workflow (FindIntegrationAnchors() and IntegrateData()), which identifies shared cell states across datasets by finding anchors between them. Clustering was based on a shared nearest neighbor (SNN) graph and dimensionality reduction via UMAP. A clustering resolution of 0.3 was selected using Clustree v0.5.1 (86). To identify cell types, reference data from Emont et al. (2022)(87) was used. Both reference and query datasets were SCT-normalized prior to label transfer via FindTransferAnchors() and TransferData() using the "cell_type__custom" column. Clusters identified as Neutrophils, Natural Killer cells and Mast cells all had less than 20 nuclei, so they were removed from subsequent analyses. Clusters identified to contain adipocytes (Seurat clusters 0, 1, 2, and 4) were sub-clustered. After re-integration and UMAP, a resolution of 0.4 was chosen (via Clustree). Adipocyte subtype labels were transferred using the "cell_subtype__custom" column from the same reference dataset. Marker identification for each major and adipocyte cluster were performed with the FindAllMarkers() function from Seurat. Likewise, differential gene expression analysis between Flox Control and PRDM16cKO samples was performed with the FindMarkers() function from Seurat. Subsequent Gene Set Enrichment analyses (GSEA) were performed using the fgsea package in R (v1.28.0) (88), with GMT files corresponding to the MSigDB C2 curated gene sets (v7.4), C3 regulatory target gene sets (v7.4), and C5 gene ontology (GO) sets (7.4).

Cryo-sectioning and Immunofluorescence

Tissues were fixed in 4% PFA at 4°C overnight and subsequently washed with 1x PBS for 1h at RT three times. Samples were then delipidated and permeabilized as described in the Adipo-Clear protocol (89). Fully delipidated samples were incubated in 25% sucrose/PBS solution overnight until sinking, and then frozen in Tissue-Tek O.C.T Compound (Sakura Finetek USA, 4583). Frozen samples were sectioned into 10μm slices using a Leica CM3050 S cryostat. Cryo-sections were placed in PBS/0.05% Triton X-100 for 1h and blocked with PBS/0.1% Triton X-100/0.05% Tween 20/2μg/ml heparin (PtxwH buffer) containing 3% donkey serum for 1h at RT, and then incubated with primary antibodies diluted in PtxwH buffer at RT overnight. Samples were then rinsed in PtxwH buffer for 5 min, 10 min, and 30 min to remove unbound antibodies. Secondary antibodies diluted in PtxwH buffer were then applied to samples at RT for 4 hr. Samples were next rinsed with PtxwH buffer for 5 min, 10 min, and 30 min, followed by 1x PBS for 10 min twice. Finally, samples were immersed in antifade mountant (ProLong Gold, ThermoFisher Scientific, P10144) and sealed with a coverslip. Anti-UCP1 (1:2000, ab10983 Abcam, RRID:AB_2241462), Anti-ALDH3aB (1:200, Proteintech #15746-1-AP, RRID:AB_2242461), Anti-COL1α1 (1:200, Cell Signaling 72026, RRID:AB_2904565), Anti-QSOX1 (1:50, Proteintech 12713-1-AP, RRID:AB_2173314) and Alexa-647 conjugated anti-rabbit secondary (1:200, Thermo Fisher Scientific, A32795, RRID:AB_2762835) antibodies were used for staining cryo-sections. Fluorescently labeled samples were imaged on an inverted LSM 980 laser scanning confocal microscope (Zeiss) with a 5X or 20X lens.

Adipo-Clear

Tissues were fixed in 4% PFA at 4°C overnight and subsequently washed with 1x PBS for 1h at RT three times. Samples were washed in 20%, 40%, 60%, 80% in $H_2O$/0.1% Triton X-100/0.3 M glycine (B1N buffer, pH 7), and 100% methanol for 30 min each followed by delipidation with 100% dichloromethane (DCM; Sigma-Aldrich) for 30 min three times. After delipidation, samples were washed in 100% methanol for 30 min twice, then in 80%, 60%, 40%, 20% methanol in B1N buffer for 30 min each step. All procedures above were carried out at 4°C with shaking. Samples were then washed in B1N for 30 min twice followed by permePBS/0.1% Triton X-100/0.05% Tween 20/2 μg/ml heparin (PTwH buffer) for 1hr twice before further staining procedures. For immunolabeling samples were incubated in primary antibody dilutions in PTxwH for 4 days. After primary antibody incubation, samples were washed in PTxwH for 5 min, 10 min, 15 min, 30 min, 1 hr, 2 hr, 4 hr, and overnight, and then incubated in secondary antibody dilutions in PTxwH for 4 days. Samples were finally washed in PTwH for 5 min, 10 min, 15 min, 30 min, 1 hr, 2 hr, 4 hr, and overnight. In this study, UCP1 (1:200, abcam, ab10983, RRID:AB_2241462) and CD68 (1:400, BioRad, MCA1957T, RRID:AB_2074849) were used. Secondary antibodies conjugated with anti-rabbit secondary Alexa-568 (Thermo Fisher Scientific Cat# A10042, RRID:AB_2534017, 1:200) and anti-rat secondary Alexa-647 (Jackson ImmunoResearch Labs Cat# 712-605-153, RRID:AB_2340694, 1:200). For tissue clearing samples were dehydrated in 25%, 50%, 75%, 100%, 100% methanol/$H_2O$ series for 30 min at each step at RT. Following dehydration, samples were washed with 100% DCM for 30 min twice, followed by an overnight clearing step in dibenzyl ether (DBE; Sigma-Aldrich). Samples were stored at RT in the dark until imaging.

3D Imaging

All whole-tissue samples were imaged on a light-sheet microscope (Ultramicroscope II, LaVision Biotec) equipped with 1.3X (used for whole-tissue views with low-magnification) and an sCMOs camera (Andor Neo). Images were acquired with the ImspectorPro software (LaVision BioTec). Samples were placed in an imaging reservoir filled with DBE and illuminated from the side by the laser light sheet. The samples were scanned with the 488, 561, and 640nm laser channels.

Image Processing

All whole-tissue images and immunofluorescence images were generated using Imaris x64 software (version 8.0.1, Bitplane). Optical slices were obtained using the "orthoslicer" tool. Optical sections pictures were generated using the "snapshot" tool.

Histology

Dissected tissues were fixed in 4% PFA overnight and transferred to 70% ethanol. Paraffin embedding, sectioning, and H&E staining was performed by the Memorial Sloan Kettering Cancer Center Laboratory of Comparative Pathology. H&E-stained sections were imaged using a wide-field fluorescence/brightfield/DIC microscope (Zeiss) at the Rockefeller University Bioimaging Resource Center. For PicrosiriusRed staining mesenteric arterial beds were fixed in 4% PFA and processed for histological inclusion in paraffin. 5µm thick tissue sections were stained with PicroSirius red (0.1% w/v) by the Histopathology Technology Platform at the Research Institute of the McGill University Health Centre. Total collagen content was measured in the perivascular area (adventitia) of all arteries in each adipose tissue from each animal by bright field analysis (magnification 40x). Perivascular fibrosis (% area) was determined as the area of the red staining in adventitia relative to the whole artery, including the lumen. Arteries were analyzed using the package ImageJ 1.44p (Wayne Rasband, NIH, USA) available in http://imagej.nih.gov/ij. To account for differences in number of arteries analyzed per animal, arteries were categorized by percent fibrosis (0-5%, 5-10%, 10-15%, …, >40%), divided by the total number of arteries analyzed for a specific mouse.

Protein isolation and western blot

Frozen tissue was stored at -80˚C was pulverized in a liquid nitrogen pre-cooled metal cell crusher and transferred into a 1.5ml Eppendorf tube. RIPA buffer supplemented with cOmplete mini protease inhibitor cocktail (Roche) and PhosSTOP (Roche) was added to the frozen tissue powder and vortex thoroughly. For full lysis samples were inverted for 15min at 4˚C using a rotator. Afterwards samples were centrifuged at 21.000g for 5min to pellet debris and protein concentration of the liquid phase was determined using Pierce BCA Protein Assay Kit (Thermo Scientific) using a dilution series of bovine serum albumin as protein standards. Proteins were diluted with 4xSDS-loading dye (BioRad) and loaded in same concentration on pre-cast polyacrylamide gels (BioRad, 4-20% Mini PROTEAN) for electrophoresis. Afterwards protein was transferred to PVDF membrane using standard wet transfer. Immunoblots were incubated with anti-αTubulin (1:1000, Cell Signaling 3873S), anti-Vinculin (1:1000, Cell Signaling 4650S), anti-UCP1 (1:1000, Abcam ab209483), AGT (1:500, IBL America #28101-S), ALDH3b2 (1:1000, Proteintech #15746-1-AP), REEP6 (1:1000, Proteintech #12088-1-AP), anti-total MLC (1:1000,

Cell Signaling 3672S), Phospho-MLC(1:500, Cell Signaling 3671T), washed 3x 15min with TBST and incubated with anti-mouse (for anti-αTubulin primary) or anti-rabbit HRP-conjugated secondary antibodies (1:5000, Jackson ImmunoResearch) and developed using Western Lightning Plus-ECL (PerkinElmer) and imaged on an autoradiographic film or using a Bio-Rad Gel Doc system.

ELISA

Protein samples were thawed on ice and ELISA used following specific kit included protocols. Following ELISA were used with PVAT and/or serum or plasma as indicated. Mouse FN1 / Fibronectin (LS-Bio Sandwich ELISA Kit - LS-F3999), Mouse COL3A1/Collagen III (LS-Bio Sandwich ELISA Kit - LS-F25411), Mouse COL4a1/Collagen IV (LS-Bio, LS-F50964-1) and Human Competition-based ANGII (RayBiotech, EIA-ANGII-1).

Flow Cytometry

Adipose tissues were collected and stored in ice cold PBS until all samples were collected. Samples were digested in HBSS w/o $Ca^{2+}$ and $Mg^{2+}$ with 0.5% BSA with 10mg/ml Collagenase D (Roche 11088882001), 20µl/ml Dispase II (Roche 04942078001) and 4µl/ml 2.5M $CaCl_2$ for 30min at 37˚C shaking at 140rpm in a waterbath. Samples were vortexed after 15 minutes of digestion and placed back at 37˚C. Digestion was stopped with equal amounts of ice-cold RPMI 1640 media (Thermo Scientific 11875093 supplemented with 10%FBS, Gibco) and filtered through 100µm cell strainer and centrifuged for 10 min at 500g at 4˚C. Aspirate supernatant and resuspend pellet in 500µl ASK Lysis buffer (Gibco A1049201) and incubated for 5min at RT. 2ml FACS Buffer was added (1xDPBS (Thermo Scientific 14190144), 0.025M HEPES Solution (Sigma-Aldrich H0887-100ML), 5% BSA (Sigma-Aldrich, A9418-100G), 0.002M EDTA(Invitrogen 15575020)) and samples centrifuged for 10 min at 500g at 4˚C. Supernatant was aspirated and cells resuspended in 1xPBS. All samples except for unstained and compensation controls were stained with live dead stain (1:1000, Zombie-Green, Biolegend 423112) for 15min at RT. For zombie-green compensation control 50µl of cell suspension was boiled for 5min at 60˚C to induce cell death and added to 50µl of live cells. Cells were spun down at 500g for 5min at 4˚C and supernatant removed. Cells were blocked with purified rat anti-mouse CD16/CD32 (1:100, BD 553142) for 15 min on ice and 2X antibody staining mix prepared in BD Brilliant Staining buffer (2X Mix: CD45-APC-Cy7 1:400 (Biolegend 103116), CD11b PerCP-Cy5.5 1:400 (Biolegend 101228), F4/80 PE 1:400 (Biolegend 123116), Ly6G APC 1:200 (Biolegend 127614), MHCII PacBlue 1:400 (Biolegend 107620), CD11c BUV737 1:50 (BD 612796), CD206 BV650 1:200 (Biolegend 141723), Tim4 PeCy7 1:200 (Biolegend 130010), SiglecFBV711, 1:200 (BD 740764) and/or B220 BV605 1:100 (Biolegend 103243), CD3 AF700 1:200 (Biolegend 100216), CD4 AF647 1:50 (Biolegend 100426), CD8a BUV737 1:50 (BD 612759), TCRyd PE-Cy7 1:200 (Biolegend 118124), NK1.1 BV711 1:50 (Biolegend 108745)) was added for 30min on ice protected from light. Cells were spin down at 500g for 5min at 4˚C and cells were resuspended in FACS buffer or fixed overnight in FACS Buffer with 0.5% Formalin and washed in FACS Buffer. Samples were filtered through blue cap flow tubes (Falcon 352235) and vortexed immediately before acquisition using BD Fortessa or BD LSRII. Data were analyzed using Flow-Jo software. Outlier test was performed using Grubbs' Test/extreme studentized deviant (ESD) method with an alpha significance set to $P<0.5$.

Cell culture

Mouse primary stromal vascular fraction (SVF) cells were obtained from adipose tissues of 5–6-week-old C57BL/6J male mice by collagenase digestion (10mg/ml collagenase D, Dispase II (dilute from 50x (1g DispaseII/8.3ml PBS)), 0.01M $CaCl_2$) and plated on collagen I-coated dishes (Corning). SVF cells from subcutaneous white adipose tissue (SQ) were grown in DMEM/F-12 GlutaMAX medium (Gibco) containing 10% FBS and 1% P/S. Once grown to confluence, differentiation and maintenance of primary adipocytes was done using DMEM/F-12 GlutaMAX medium containing 10% FBS and 1% P/S. SQ SVF cells were induced to differentiate with an adipogenic cocktail (0.5 mM IBMX, 1 µM dexamethasone, 1 µM rosiglitazone, and 850 nM insulin) for the first 2 days, followed by 2 days of 1 µM rosiglitazone and 850 nM insulin, after which the cells were maintained in 850 nM insulin for additional 2days. Primary SVF and adipocytes were maintained at 37 °C with 10% $CO_2$. All cultured primary adipocytes were checked for lipid accumulation under a phase-contrast microscope during differentiation. For collection of conditioned media, adipocytes were provided with 1ml of fresh media and re-collected after 24h. Media was filtered to remove potential cell debris and frozen in liquid nitrogen. For media shuffling experiments, media was mixed in a ratio of 1:1 with fresh vascular smooth muscle media (DMEM+5%FBS+1P P/S) and added for 48h. VSMC were washed 3 times with PBS and collected and lyzed using TRIzol (Invitrogen) and a cell scraper to perform RNA isolation. For primary vascular smooth muscle isolation from mouse aorta (6-8 weeks old C57BL/6J male mice), PVAT was removed and aortas from 2 mice each pooled for one replicate. Aortas were placed in digestion buffer (1mg/ml collagenase Type II (Worthington), 1mg/ml soybean trypsin inhibitor (Worthington), 0.744units/ml elastase (Worthington) and 1%P/S in HBSS) for 8-10 minutes and adventitia was removed. Aortas were cut open and residual blood and endothelial layer mechanically removed. Briefly wash aortas in DMEM/F12 (20% FCS, 1%P/S) and place for 1h in fresh digestion buffer. Mechanically disrupt VSMC layer, add DMEM/F12 (20% FCS, 1%P/S) and centrifuge to pellet cells. Wash pellet and centrifuge again, before reconstituting in 1ml media and distribute in TC-treated 24-weel plate. Replace media after 1 week and remove all cell debris and grown to ~95% confluency before splitting (PBS wash, 5min Trypsin (Gibco)). VSMC were expanded in 24-weel plates before final seeding in 6-Well plate for experiments.

RNA isolation, cDNA synthesis, and RT-qPCR

Total RNA was extracted from cultured cells and adipose tissue by TRIzol (Invitrogen). Cells were collected by scaping them with a cell scraper in TRIzol and frozen tissue was snap frozen in liquid nitrogen and stored at -80˚C before pulverization in a liquid nitrogen pre-cooled metal cell crusher. Tissue powder was mixed with TRIZol and incubated under constant rotation for 15min at 4˚C for full lysis and centrifuged to pellet cell debris. Supernatant was used to purify total RNA using chloroform, ethanol and RNeasy Micro Kit (Qiagen). cDNA was synthesized from 1 µg of RNA using the High-Capacity cDNA Reverse Transcription Kit (Applied Biosciences). Power SYBR Green (Life Technologies) was used for RT-qPCR reactions performed with QuantStudio 6 Flex Real-Time PCR System (Thermo Scientific) in a 384-well format. Relative fold changes of mRNA levels were calculated using the ΔΔCT method with RPL as loading control. Outlier test was performed using Grubbs' Test/extreme studentized deviant (ESD) method with an alpha significance set to $P<0.5$. qPCR primers are provided in Table S2.

Human data analysis

A propensity score – matched (PSM) cohort of patients with and without detectable BAT by FDG-PET/CT scans was established as previously described (15). This cohort is comprised of 14,923 individuals (5,070 BAT positive and 9,853 BAT negative) matched by sex, age, BMI, and outside temperature at the time of their index scan (first scan that reported BAT in BAT positive patients, first scan chronologically in BAT negative patients). We then extracted echocardiogram reports up to 1 year after the index scan from the EHR of each patient belonging to the PSM cohort. Out of 14,923 patients, 1,748 had available echocardiogram reports for further analysis (1,095 BAT positive and 653 BAT negative patients). Data on LV mass index, LVPWD, and LA volume index was extracted from these reports and compared between BAT positive and BAT negative patients using a general linear model (GLM), and least square means (LSMeans) were estimated adjusting for age, sex, BMI, race, and outdoor temperature. Pairwise comparisons were performed using the least significant difference (LSD) test, and 95% confidence intervals were reported. Statistical analyses were conducted using SAS software (SAS Institute, Cary, NC, USA). This study followed institutional guidelines and was approved by the Institutional Review Boards of Memorial Sloan Kettering Cancer Center (IRB## 17-581). Due to the retrospective nature of this study, the requirement for informed consent was waived. Results from this study are reported in accordance with the Strengthening the Reporting of Observational Studies in Epidemiology guidelines for case–control studies. Report on age, sex, and ethnicity was obtained for all participants and included as a variable for the analyses.

<u>Cardiometabolic phenome-wide association studies in UKBB and MSM</u>

Cardiometabolic associations of variants in exon 9 of *PRDM16* were investigated in the UKBB and MSM, comprising exome sequencing (ES) data and electronic health records (EHR) of 189,448 and 58,990 participants, respectively. Genetic ancestry was determined based on genetic similarity to reference populations from 1000 Genomes Project using ADMIXTURE (90,91). Participants with EUR ancestry in UKBB (n = 180,500) and MSM ($n$ = 21,680), as well as AFR ancestry in MSM ($n$ = 19,856) were included in the analyses. Patient diagnoses were extracted from EHR based on ICD-9 and ICD-10 codes, which were then mapped to phecodeX (92). 386 phecodes (binary phenotypes) from the "Cardiovascular" and "Endocrine/Metabolic" categories, along with nine quantitative measurements i.e., systolic and diastolic blood pressure, BMI, total cholesterol, LDL, HDL, triglycerides, blood glucose and HbA1c were extracted from EHR.

For all three cohorts, a sparse genetic relationship matrix was constructed using 5,000 linkage disequilibrium (LD)-pruned random markers with a relatedness cut-off of 0.125 (93,94). The null model for each phenotype was fitted using age, sex and first ten genetic principal components as covariates. Protein altering variants located in exon 9 of *PRDM16* were extracted, resulting in 258 missense variants in UKBB and 206 missense variants in MSM respectively. Pairwise LD was calculated with an $r^2$ threshold of 0.2, and independent variants with a minor allele count greater than 20 were selected, which identified 25 variants in UKBB, 7 variants in MSM-EUR and 12 variants in MSM-AFR. Association testing was conducted using SAIGE for binary phenotypes and inverse normalized quantitative traits (94). *P* values were adjusted using the Benjamini-Hochberg method.

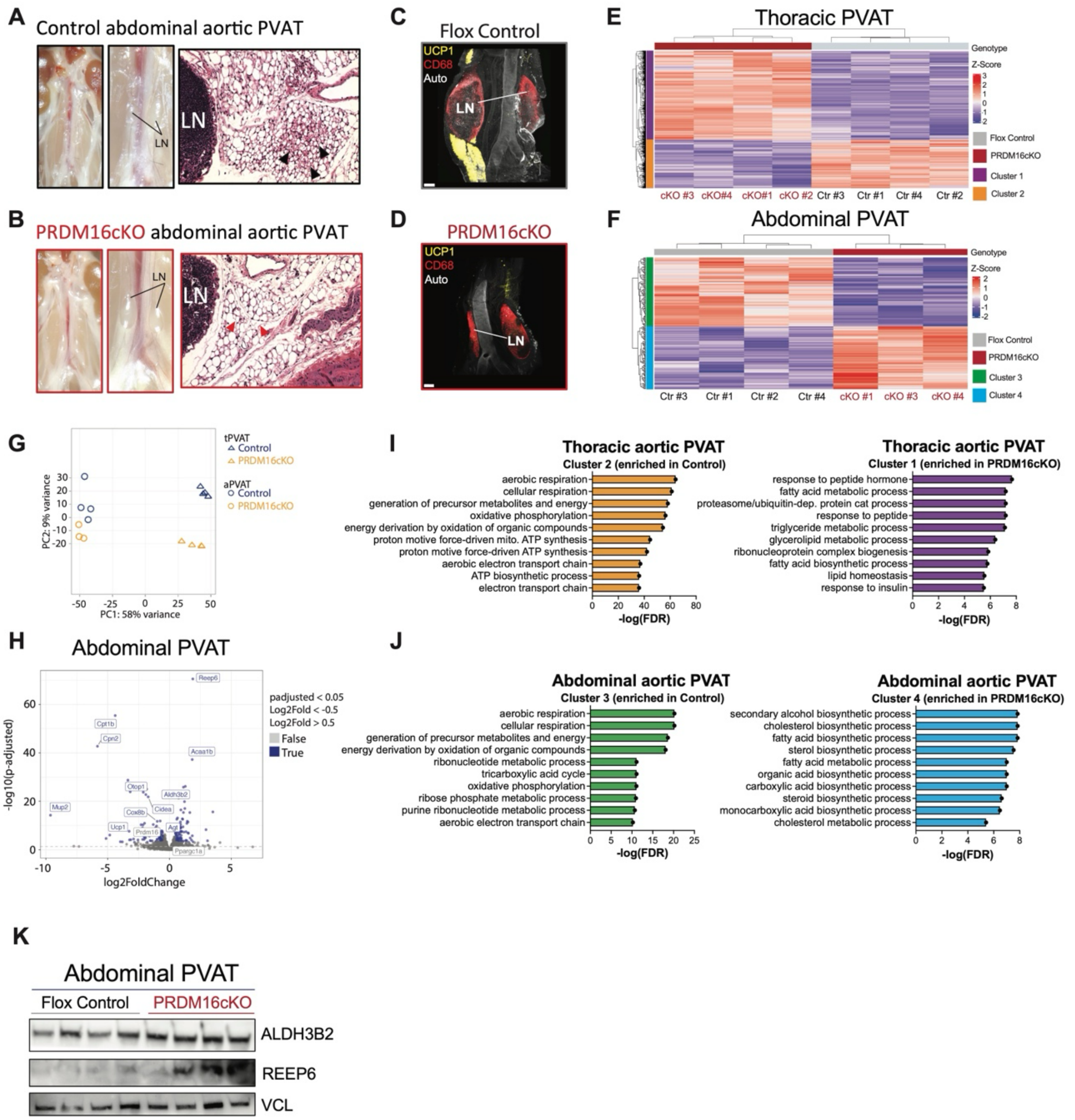

**Fig. S1 Loss of *Prdm16* in adipocytes leads to remodeling of abdominal and thoracic aortic PVAT.** (**A, B**) Representative image and HE stain of the thoracic PVAT in (A) PRDM16flox control (flox control) and (B) *Prdm16*flox; *AdipoQ*-Cre+ (PRDM16cKO) mice. (**C, D**) Whole mount immunolabeling with UCP1 and CD68 and tissue clearing of aPVAT from (C) flox control and (D) PRDM16cKO mice, green laser was used for visualization of aorta and surrounding tissue by autofluorescence. Scale bars are 200µm. (**E, F**) Heatmap of differentially expressed genes (DEGs) between PRDM16cKO (n=3-4) and flox control (n=4) (E) thoracic PVAT and (F) abdominal PVAT. (**G**) Principal component analysis (PCA) plot of tPVAT and aPVAT of PRDM16cKO (n=3-4 per group) and flox control mice (n= 4 per group). Individual datapoints represents data from a single animal. (H) Volcano plot for DEGs in the abdominal PVAT of PRDM16cKO minus flox control; blue color highlights significantly up-/down-regulated genes (padj. <0.05) and $|log_2[\text{fold change}]| \geq 0.5$ and grey color display non-significant DEGs. Some significant DEGs related to thermogenesis and most upregulated in PRDM16cKO are highlighted.

(**I, J**) Pathway enrichment analysis showing the 10 most enriched pathways in (I) tPVAT (yellow, enriched in flox control, purple enriched in PRDM16cKO) and (J) aPVAT (green, enriched in flox control, blue enriched in PRDM16cKO). (**K**) Representative western blots of ALDH3B2, REEP6 and VINCULIN (VCL) of flox control (n=4) and PRDM16cKO (n=4) abdominal PVAT.

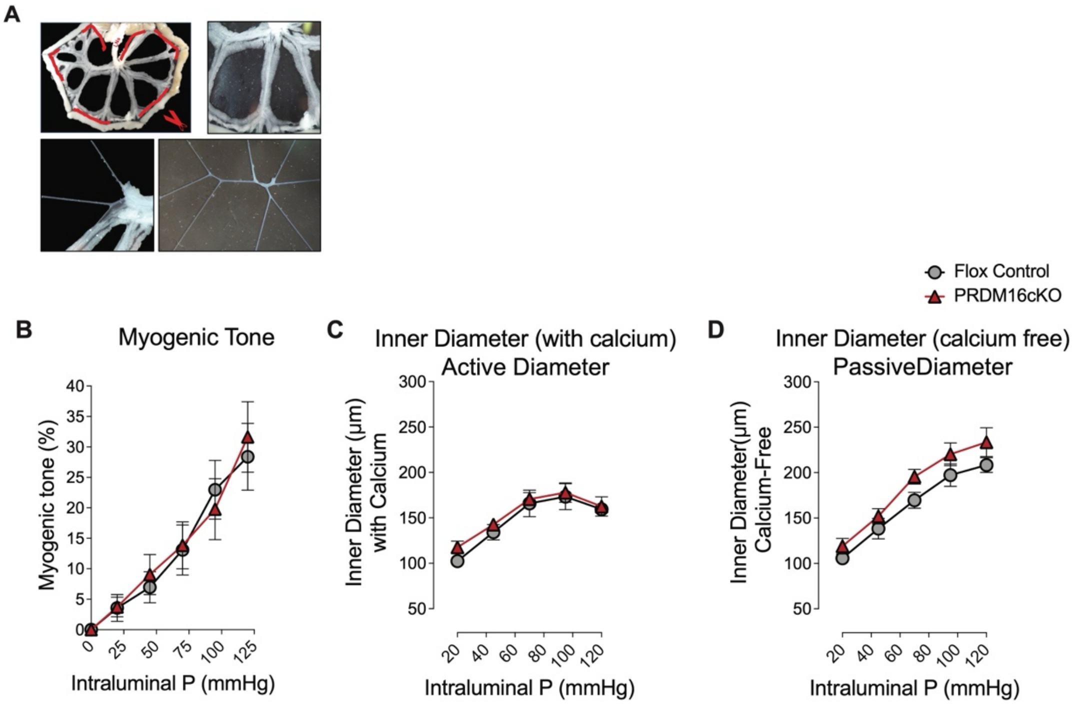

**Fig. S2 Ablation of beige fat does not affect myogenic tone in PRDM16cKO mice.** (**A**) Representative image of mesenteric adipose tissue and artery isolation for pressure myograph and single nuclear RNA sequencing experiments. Red dotted line represents cuts to remove mesenteric tissue from the gut and the mesenteric lymph nodes (LN) (**B-C**) Pressure myography of mesenteric arteries after removal of perivascular fat was conducted in flox control and PRDM16cKO mice (n=4-5 per group; 1-2 mesenteric arteries per mouse). (B) Myogenic tone was calculated as a function of ((passive diameter (D)) – active diameter (C))/(passive diameter (D)))*100. Data are mean+/- SEM.

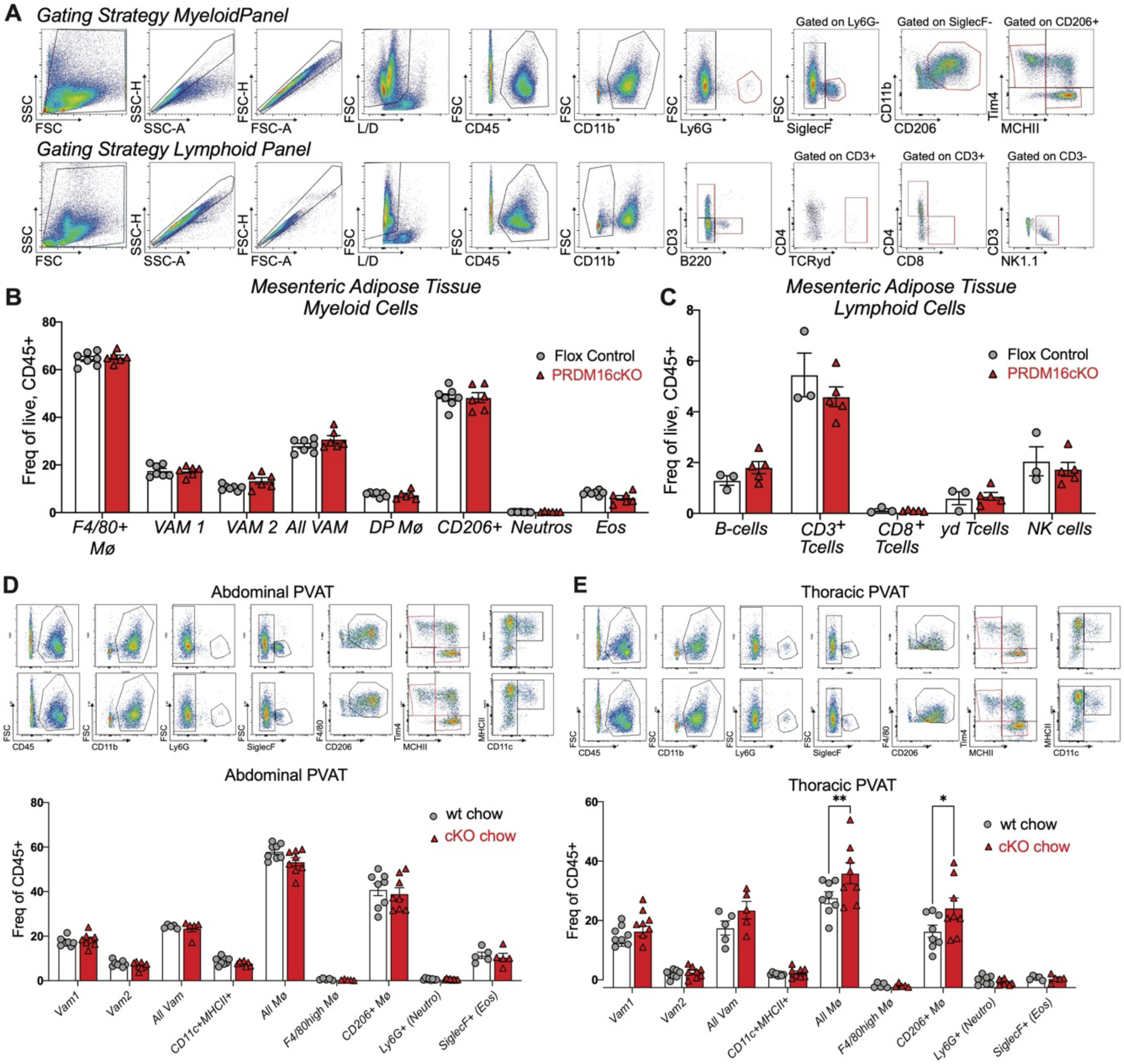

**Fig. S3 Flow Cytometry of PVAT depots did not show changes in immune cell composition between PRDM16cKO and flox control mice.** (**A**) Representative gating strategy of mPVAT (top panel myeloid gating, bottom panel lymphoid panel). (**B, C**) Flow cytometry analysis of mPVAT for (B) myeloid and (C) lymphoid immune cell populations of PRDM16cKO (n=5-6) and flox control mice (n=3-7). Individual data points represent data from a single animal, and bars are means ± SEMs. Significance was calculated using two-way analysis of variance (ANOVA), with **P* < 0.05; ***P* < 0.01; *****P* < 0.0001; ns, not significant. (**D, E**) Representative gating strategy (top) and flow cytometry analysis of (D) abdominal aortic PVAT and (E) thoracic aortic PVAT of PRDM16cKO (n=8) and flox control mice (n=8). Two animals were pooled during tissue digestion for each individual data point and data represent results from two independently experiments. Bars are means ± SEMs. Significance was calculated using two-way analysis of variance (ANOVA), with **P* < 0.05; ***P* < 0.01; *****P* < 0.0001; ns, not significant.

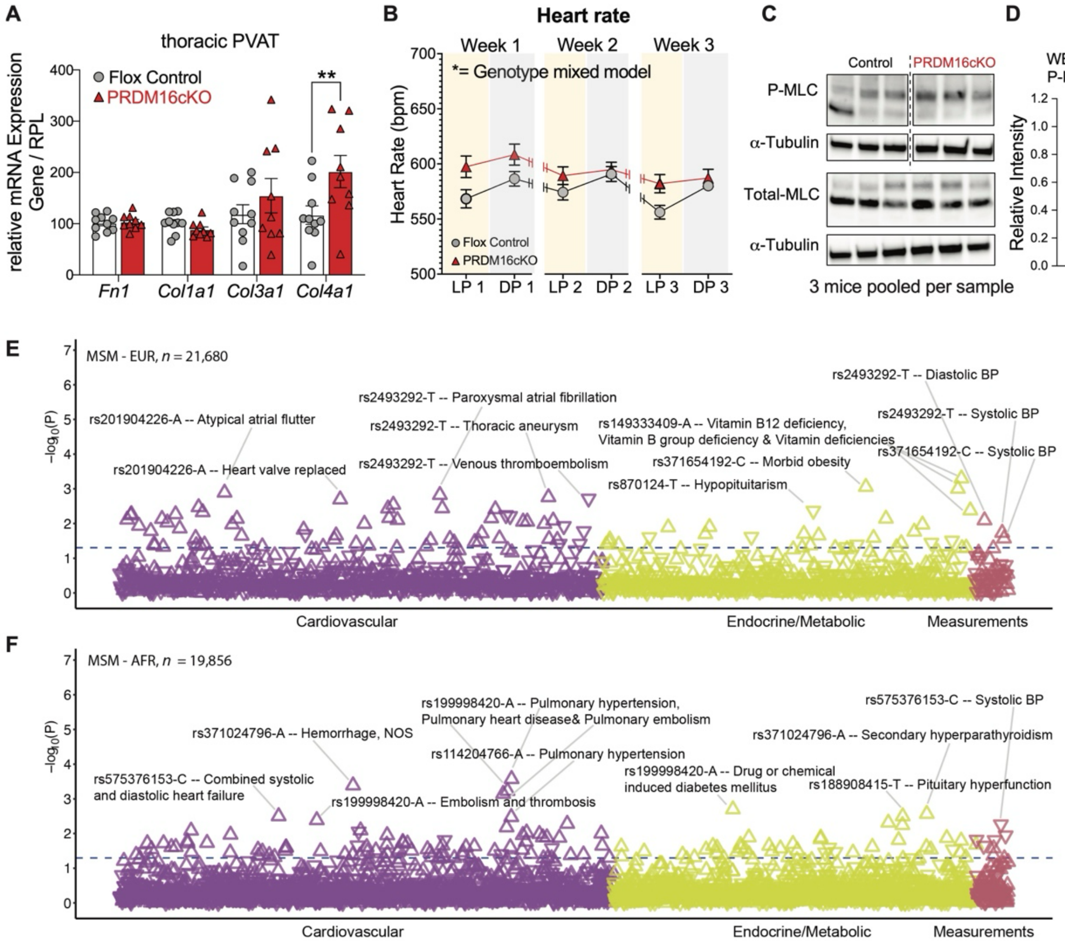

**Fig.S4 Loss of *Prdm16* in adipocytes leads to increased extracellular matrix expression and elevated blood pressure.** (**A**) qPCR for *Fn1*, *Col1a1*, *Col3a1* and *Col4a1* on thoracic aortic PVAT of flox control (n=10) and PRDM16cKO (n=9) mice. Individual data points represent data from a single animal, and bars are means ± SEMs. Significance was calculated using unpaired Student's *t* test. *$P < 0.05$; **$P < 0.01$; ****$P < 0.0001$; ns, not significant. (**B**) Measurements of heart rate recorded with implanted radiotelemetry devices in freely moving mice (n=28-29 per group combined from 3 independent experiments). Data are mean+/-SEM. A linear mixed model for repeated measures over time was used to analyze the radiotelemetry data. *$P < 0.05$; **$P < 0.01$; ****$P < 0.0001$; ns, not significant. (**C**) Western blot for total and phosphorylated (P-) myosin light chain (MLC) and αTUBULIN on mesenteric arteries. A total of 5 mesenteric arteries of 3 mice were pooled in each sample. (**D**) Quantification of the western blot by ImageJ. (**E, F**) Ancestry-specific analysis of associations of exon 9 variants with cardiovascular and endocrine/metabolic traits in the European (EUR) (E) and African (F) Mount Sinai Million (MSM) cohorts. The direction of the triangles indicates the direction of effect (upward: increased risk or level, downward: decreased risk or level). The blue dashed line represents $P = 0.05$.

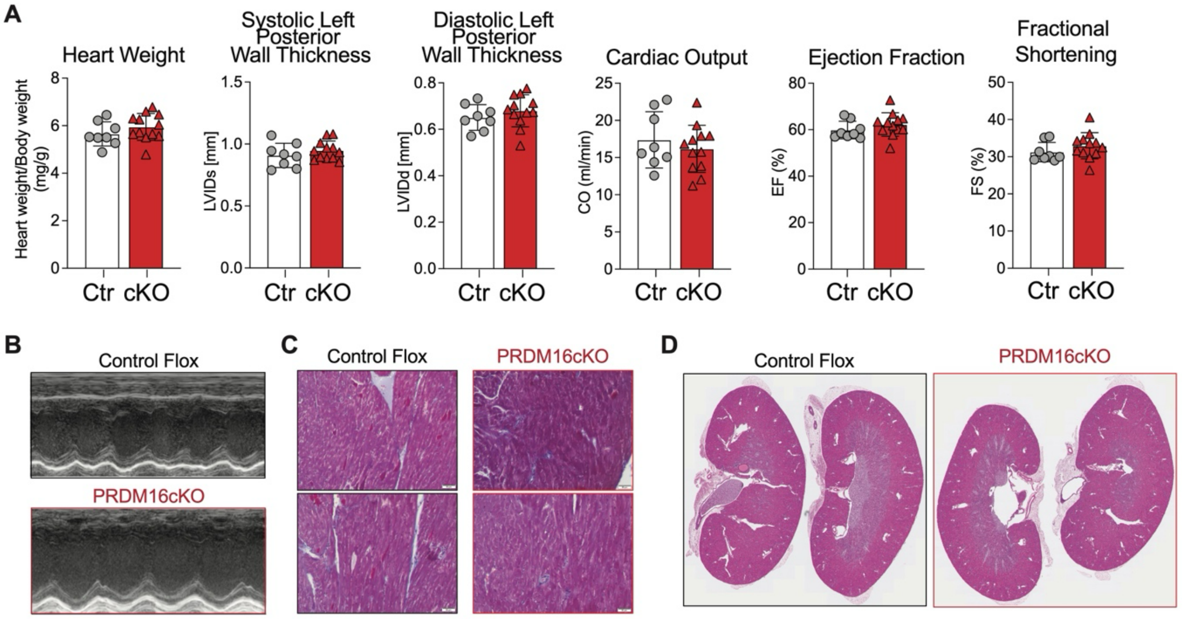

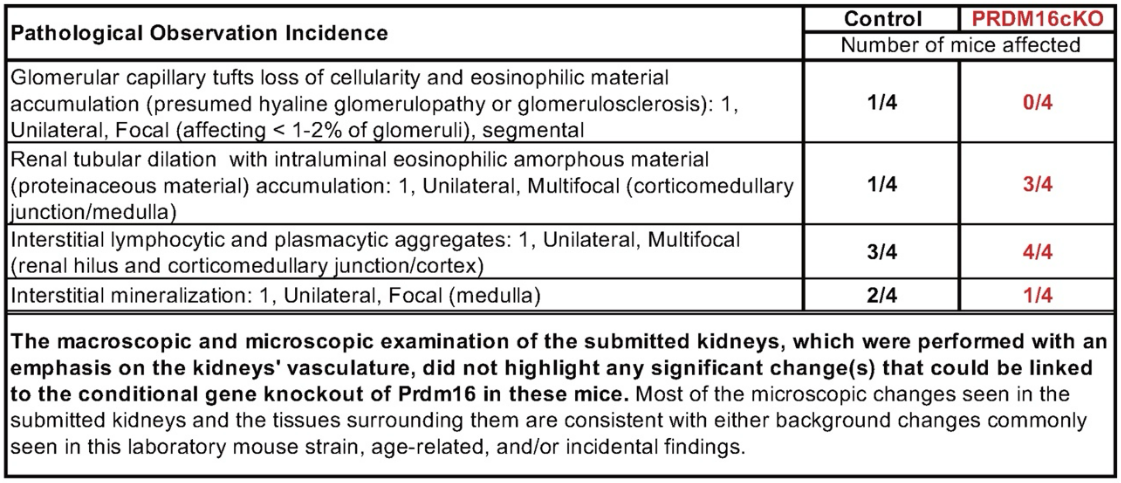

E

**Pathological Assesment Kidney - Focus on Vasculature**

| Pathological Observation Incidence | Control | PRDM16cKO |
|---|---|---|
| | Number of mice affected | |
| Glomerular capillary tufts loss of cellularity and eosinophilic material accumulation (presumed hyaline glomerulopathy or glomerulosclerosis): 1, Unilateral, Focal (affecting < 1-2% of glomeruli), segmental | **1/4** | **0/4** |
| Renal tubular dilation with intraluminal eosinophilic amorphous material (proteinaceous material) accumulation: 1, Unilateral, Multifocal (corticomedullary junction/medulla) | **1/4** | **3/4** |
| Interstitial lymphocytic and plasmacytic aggregates: 1, Unilateral, Multifocal (renal hilus and corticomedullary junction/cortex) | **3/4** | **4/4** |
| Interstitial mineralization: 1, Unilateral, Focal (medulla) | **2/4** | **1/4** |

**The macroscopic and microscopic examination of the submitted kidneys, which were performed with an emphasis on the kidneys' vasculature, did not highlight any significant change(s) that could be linked to the conditional gene knockout of Prdm16 in these mice.** Most of the microscopic changes seen in the submitted kidneys and the tissues surrounding them are consistent with either background changes commonly seen in this laboratory mouse strain, age-related, and/or incidental findings.

**Fig. S5 No changes in heart function or pathology of heart and kidney in PRDM16cKO mice.** (**A, B**) Echocardiography (A) results and (B) representative ultrasound images of PRDM16cKO (n=12) and flox control (n=8) mice. Individual data points represent data from a single animal, and bars are means ± SEMs. (**C**) Representative images of Masson's trichrome staining of heart sections. Images show the left ventricular wall of PRDM16cKO and flox control mice. Scale bar is 50µm. (**D**) Representative images of haematoxylin/eosin (HE) staining of kidneys from PRDM16cKO and flox control mice and (**E**) pathological assessment of the kidney with focus on the vasculature. Focal (for glomerular changes: involving 50% of glomeruli).

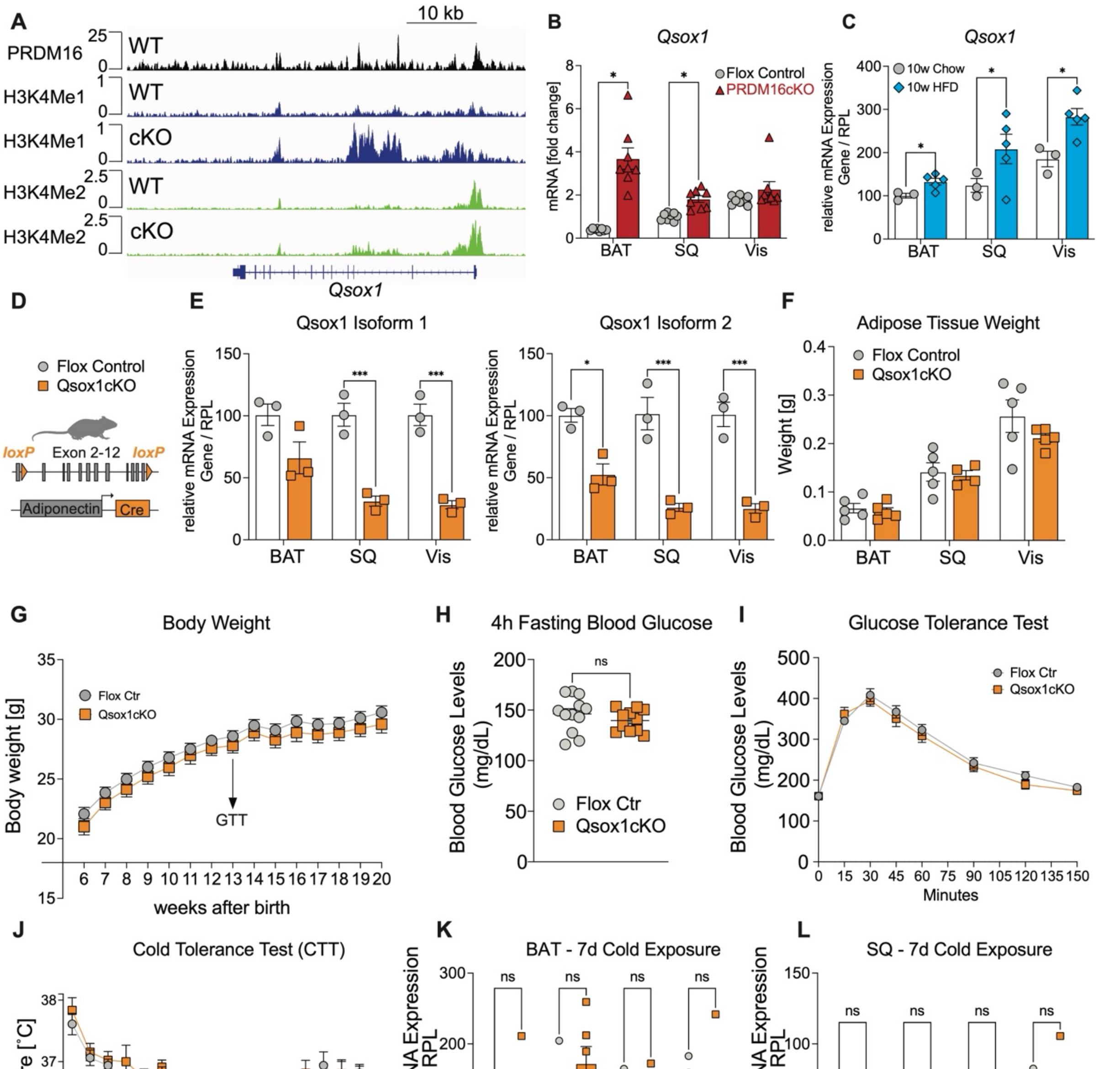

**Fig. S6 Loss of *Qsox1* in adipocytes does not affect whole body metabolism.** (**A**) Analysis of published PRDM16 ChIP sequencing data (39) for binding of PRDM16 on *Qsox1* in control and PRDM16cKO thermogenic brown adipose tissue (**B-C**) qPCR analysis of *Qsox1* in brown (BAT), subcutaneous (SQ) and visceral (Vis) adipose tissue from (B) PRDM16cKO and flox control mice and (C) control mice fed a standard (chow) or high-fat diet (HFD) for 10 weeks. (**D**) Schematic of *Qsox1*$^{loxP/loxP}$; *AdipoQ*-Cre-/+ (Qsox1cKO) mice. (**E**) qPCR of Qsox1 (Isoform1 and 2) in BAT, SQ and Vis adipose tissue from flox control and Qsox1cKO mice (n=3 per group). (**F**) Adipose tissue weights form BAT, SQ and Vis adipose tissue of Qsox1cKO and flox control (n=4-5) mice.

(**G**) Body weight of Qsox1cKO (n=11) and flox control mice (n=12) over a period of 14 weeks. Data are mean+/- SEM. (**H**) 4h fasting blood glucose and (**I**) intraperitoneal glucose tolerance test (2g/kg) of Qsox1cKO (n=11) and flox control mice (n=12) at 13 weeks of age. Data in (I) are mean+/- SEM. Statistical analysis in (I) two-way analysis of variance (ANOVA) for repeated measure with Šídák correction for multiple comparisons, with $*P < 0.05$; $**P < 0.01$; $****P < 0.0001$; ns, not significant. (**J**) Cold tolerance test of Qsox1cKO (n=9) and flox control (n=8) mice over a 7-day period. (**K, L**) qPCR of thermogenic genes (*Ucp1*, *PGC1a*, *Cidea*, *Elovl3*) in (K) BAT and (L) SQ adipose tissue from 7 day cold-exposed flox control and Qsox1cKO mice (n=6 per group). Individual data points in (B, C, E, F, H, K, L) represent data from a single animal, and bars are means ± SEMs. Significance for (B, H) was calculated using Student's *t* test with multiple comparison correction $*P < 0.05$; $**P < 0.01$; $****P < 0.0001$; ns, not significant. Significance for (C, E, K and L) was calculated using a two-way analysis of variance (ANOVA), with $*P < 0.05$; $**P < 0.01$; $****P < 0.0001$; ns, not significant.

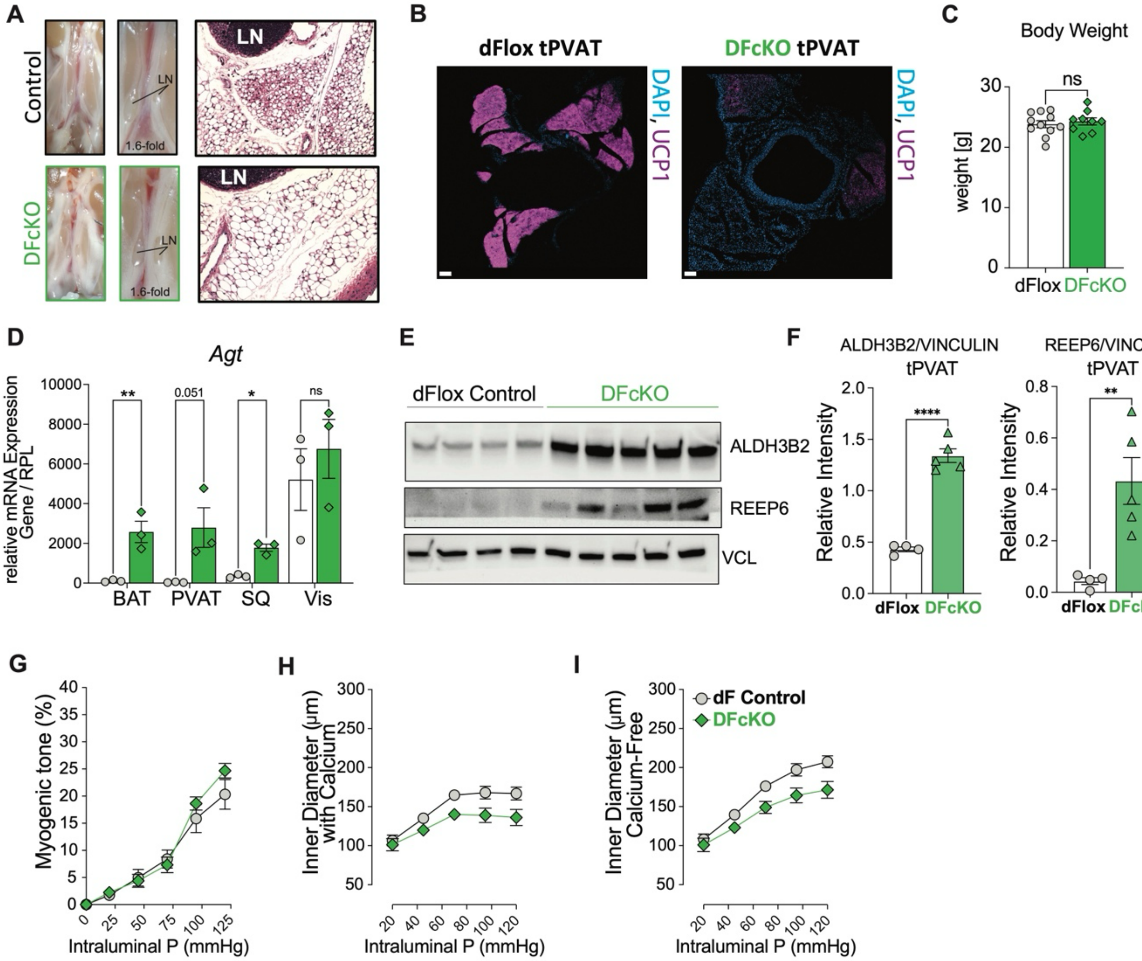

**Fig. S7 Thermogenesis in DFcKO mice is impaired.** (**A**) Representative image and HE stain of the abdominal PVAT in *Qsox1*$^{loxP/loxP}$; *Prdm16*$^{loxP/loxP}$ (dFlox control, grey) mice and *Qsox1*$^{loxP/loxP}$; *Prdm16*$^{loxP/loxP}$; *AdipoQ*-Cre-/+ (DFcKO, green) mice. (**B**) Representative immunofluorescence image for UCP1 and DAPI nuclei staining on tPVAT. Scale bar 100µm. (**C**) Body weight of DFcKO (n=9) and dFlox control mice (n=11) with 6 weeks of age. Data are mean+/- SEM. Statistical analysis was performed using Student's *t* Test, with **P* < 0.05; ***P* < 0.01; *****P* < 0.0001; ns, not significant. (**D**) qPCR of adipose tissue for *Agt* on brown (BAT), thoracic aortic PVAT (tPVAT), subcutaneous (SQ) and visceral (Vis) adipose tissue from dflox control and DFcKO mice (n=3 per group). (**E**) Representative western blots of ALDH3b2, REEP6 and VINCULIN (VCL) of dflox control (n=4) and DFcKO (n=4) tPVAT and (**F**) quantification. (**G-I**) Pressure myography of mesenteric arteries after removal of perivascular fat was conducted in dflox control and DFcKO mice (n=5 per group; 1-2 mesenteric arteries per mouse). (G) Myogenic tone calculated from (H) inner diameter with and (I) without calcium. Individual data points in (C, D and F) represent data from a single animal, and bars are means ± SEMs. Significance for was calculated using Student's *t* test **P* < 0.05; ***P* < 0.01; *****P* < 0.0001; ns, not significant.

**Supplement Table 1.** Linear mixed models for repeated measures analysis of radiotelemetry data adjusted for heart rate.

| | | |
|---|---|---|
| **Systolic Blood Pressure** | | |
| **Type III Tests of Fixed Effects** | | |
| Independent Variable | Num DF | Den DF |
| Genotype (KO) | **1** | **66** |
| Week | 2 | 66 |
| Genotype*Week | 2 | 66 |
| HR (per bpm) | **1** | **66** |
| **Estimated Mean Differences for KO vs. WT** | | |
| Week | Estimate | 95% CI Lower |
| 1 | 1.70 | -1.48 |
| 2 | 3.76 | 0.42 |
| 3 | 3.04 | -0.02 |
| Over all 3 weeks | **2.83** | **0.17** |
| **Diastolic Blood Pressure** | | |
| **Type III Tests of Fixed Effects** | | |
| Independent Variable | Num DF | Den DF |
| Genotype (KO) | **1** | **66** |
| Week | 2 | 66 |
| Genotype*Week | 2 | 66 |
| HR (per bpm) | **1** | **66** |
| **Estimated Mean Differences for KO vs. WT** | | |
| Week | Estimate | 95% CI Lower |
| 1 | 1.84 | -1.10 |
| 2 | 3.80 | 0.22 |
| 3 | **4.18** | **0.80** |
| Over all 3 weeks | **3.27** | **0.36** |
| **Mean Arterial Pressure** | | |
| **Type III Tests of Fixed Effects** | | |
| Independent Variable | Num DF | Den DF |
| Genotype (KO) | **1** | **66** |
| Week | 2 | 66 |
| Genotype*Week | 2 | 66 |
| HR (per bpm) | **1** | **66** |
| **Estimated Mean Differences for KO vs. WT** | | |
| Week | Estimate | 95% CI Lower |
| 1 | 2.13 | -0.42 |
| 2 | **3.91** | **1.00** |
| 3 | **3.91** | **1.24** |
| Over all 3 weeks | **3.31** | **1.02** |

The Bonferroni adjusted significance level for post-hoc testing at weeks 1, 2, and 3 is 0.0167.

**Table S1.** Linear mixed models for repeated measures analysis of radiotelemetry data adjusted for heart rate.

### Supplement Table 2

Primer Sequences qRT-PCR
Manuscript: Koenen M et al. 2025

| Name | strand | direction | Sequence | |
|---|---|---|---|---|
| Acta2 | for | 5' | AGCCATCTTTCATTGGGATGG | 3' |
| | rev | 5' | CCCCTGACAGGACGTTGTTA | 3' |
| AGT | for | 5' | GTCTGTGCCCATGATCTCCGGC | 3' |
| | rev | 5' | GCACGCACGTCACGGAGAAGT | 3' |
| AT1a | for | 5' | AACAGCTTGGTGGTGATCGTC | 3' |
| | rev | 5' | CATAGCGGTATAGACAGCCCA | 3' |
| AT1b | for | 5' | TGGCTTGGCTAGTTTGCCG | 3' |
| | rev | 5' | ACCCAGTCCAATGGGGAGT | 3' |
| AT2 | for | 5' | AGCCTGCATTTTAAGGAGTGC | 3' |
| | rev | 5' | AAGGACGGCTGCTGGTAATG | 3' |
| Cidea | for | 5' | GCCGTGTTAAGGAATCTGCTG | 3' |
| | rev | 5' | TGCTCTTCTGTATCGCCCAGT | 3' |
| Col1a1 | for | 5' | GCTCCTCTTAGGGGCCACT | 3' |
| | rev | 5' | CCACGTCTCACCATTGGGG | 3' |
| Col3a1 | for | 5' | CTGTAACATGGAAACTGGGGAAA | 3' |
| | rev | 5' | CCATAGCTGAACTGAAAACCACC | 3' |
| Col4a1 | for | 5' | TCCGGGAGAGATTGGTTTCC | 3' |
| | rev | 5' | CTGGCCTATAAGCCCTGGT | 3' |
| Elovl3 | for | 5' | TCC GCG TTC TCA TGT AGG TCT | 3' |
| | rev | 5' | GGA CCT GAT GCA ACC CTA TGA | 3' |
| Fn1 | for | 5' | GATGTCCGAACAGCTATTTACCA | 3' |
| | rev | 5' | CCTTGCGACTTCAGCCACT | 3' |
| PGC1a | for | 5' | CCCTGCCATTGTTAAGACC | 3' |
| | rev | 5' | TGCTGCTGTTCCTGTTTTC | 3' |
| PRDM16 | for | 5' | CAGCAGGGTGAAGCCATTC | 3' |
| | rev | 5' | GCGTGCATCCGCTTGTG | 3' |
| Qsox1 Iso 1 | for | 5' | CTCTCAGGTGCTCTGAGTGAGG | 3' |
| | rev | 5' | TACATCTAGGGCAGTGGCTCC | 3' |
| Qsox1 Iso 2 | for | 5' | CCGAGCTGCTCTTGTGAAGT | 3' |
| | rev | 5' | CTCTGGACTTGTCTGCCTCAA | 3' |
| Ucp1 | for | 5' | CTTTGCCTCACTCAGGATTGG | 3' |
| | rev | 5' | ACTGCCACACCTCCAGTCATT | 3' |
| Rpl | for | 5' | CCTGCTGCTCTCAAGGTT | 3' |
| | rev | 5' | TGGCTGTCACTGCCTGGTACTT | 3' |

**Table S2.** Primer Sequences qRT-PCR
